# *JAG2*-related muscular dystrophy and Notch signaling dysfunction in muscle stem cells

**DOI:** 10.1101/2025.07.23.665646

**Authors:** Nam Chul Kim, Minoru Tanaka, Isabelle Draper, Hannah R. Littel, Mekala Gunasekaran, Johnnie Turner, Natalya M. Wells, Qasim Mujteba, Yoko Asakura, Peter B. Kang, Atsushi Asakura

## Abstract

We previously identified a muscular dystrophy caused by biallelic variants in *JAG2*, whose protein product Jagged2 is a canonical Notch ligand. However, the disease mechanism remains unclear, particularly with respect to muscle stem cell (MuSC) function and muscle regeneration. We examined the consequences of *JAG2* deficiency and modeled pathogenic *JAG2* variants *in vitro* and *in vivo*, the latter in mouse and fly models and with particular attention to the MuSC-muscle endothelial cell (MuEC) niche. We found that both *Jag2* deficiency and overexpression of pathogenic *JAG2* variants impaired Notch signaling and myogenic self-renewal and differentiation. Hypomorphic *Jag2* mutant (*Jag2^sm^*) mice display depleted MuSCs, corresponding with impaired muscle regeneration in those mice. Co-culture experiments and the examination of cell-type-specific *Jag2* conditional knockout mice demonstrated that MuEC-specific *Jag2* knockout resulted in reduced MuSC self-renewal, while MuSC-specific *Jag2* knockout resulted in reduced myogenic differentiation. Human reference *JAG2*, but not human pathogenic variants of *JAG2*, rescued the deficiency of *Serrate (Ser)*, the *Drosophila* ortholog of *JAG2*. Therefore, pathogenic variants in *JAG2* impair muscle development and regeneration through disrupted cell-autonomous *cis-* inhibition and non-autonomous *trans-*activation involving Notch signaling dysfunction. Our findings indicate that optimizing JAG2-mediated Notch signaling is a potential therapeutic approach for *JAG2*-related muscular dystrophy.

## INTRODUCTION

Muscle stem cells (MuSCs), also known as satellite cells, are normally quiescent cells located underneath the basal lamina of muscle fibers. In response to injury, MuSCs activate, proliferate, differentiate, and either form new myofibers or fuse with existing myofibers to repair the damaged muscle. A small proportion of activated MuSCs self-renew or escape activation in order to replenish the MuSC pool(1–4). Defects in self-renewal lead to fewer MuSCs and diminished muscle regenerative capacity, particularly in aged and diseased muscle(1, 5–7). Prior studies have elucidated the molecular mechanisms regulating MuSCs through the Notch, Wnt, FGF, extracellular matrix signaling pathways, as well as juxtacrine interactions(8–10). We and others have studied the role of muscle endothelial cells (MuECs) in regulating MuSCs(11–14). We have shown that increasing vascular density can augment MuSC numbers(15–17), which is mediated through the activation of Notch signaling in MuSCs by Dll4, an MuEC-derived Notch ligand(14).

In mammalian cells, Notch signaling involves the transmembrane Notch ligands Jag1, Jag2, Dll1, Dll3, and Dll4, which bind to Notch receptors 1-4(18, 19), and has been implicated in the homeostasis of stem cells, including MuSCs. Notch signaling regulates the maintenance of MuSC quiescence(9, 20–22) as well as the formation of the MuSC niche (23). However, the precise mechanisms of Notch ligand contributions to niche formation and Notch activation in MuSCs remain unclear. Endothelial cells express Notch ligands, which in turn regulate stem cells(24), including hematopoietic stem cells (HSC)(25–28), neural stem cells(29), and MuSCs(14).

Disruption of Notch signaling is associated with several skeletal muscle diseases, particularly those associated with *JAG2*(30), *MEGF10*(31, 32), *POGLUT1*(33), and *NOTCH2NLC*(34). We discovered that biallelic pathogenic variants in *JAG2* are associated with congenital muscular dystrophy (CMD) and limb-girdle muscular dystrophy (LGMD)(30). Human JAG2 is a 1,238 amino acid membrane protein that interacts with Notch receptors via extracellular epidermal growth factor-like (EGF) domains(35), triggering cell-cell interaction-mediated *trans-* activation of Notch signaling(36). JAG2 binding to a Notch receptor leads to double cleavage, followed by migration of the Notch intracellular domain (NICD) to the nucleus, where it regulates transcription(37, 38).

The orthologous *Serrate* gene in *Drosophila* was identified in 1990(39), followed by *Jag2* in rats(40), *Jag2* in mice(41), and *JAG2* in humans(41–43). JAG2 is expressed in mammalian skeletal muscle(44), brain(45, 46), gastrointestinal tract(47), including the enteric nervous system(48), immune system(49, 50), ovarian follicles(51, 52), and endothelial cells(14, 53, 54). JAG2 is expressed in MuECs and MuSCs(14).

Notch ligands suppress or activate Notch signaling cell-autonomously, through *cis-* inhibition(55–58) or *cis-*activation(59), or via *trans-*activation of Notch receptors on neighboring cells. The exact ligand-receptor mechanisms that regulate Notch signaling in MuSCs remain unclear. For the current study, we examined the *cis-* and *trans-*regulatory activities of Jag2 for MuSC function and skeletal muscle regeneration in *Jag2* hypomorphic (*Jag2^sm^*) mice, as well as in conditional MuEC-specific and MuSC-specific *Jag2* knockout mice. We utilized MuEC-MuSC coculture systems to examine Jagged2-mediated *trans-*activities of Notch for MuSCs. Lastly, we introduced reference and variant forms of human *JAG2* into MuSC cultures and *Drosophila*.

## RESULTS

### Jag2 expression patterns in MuSCs and MuECs

Jag2-Notch signaling is important for cell communication in skeletal muscle(60–62). Previously, we performed a directional interactome analysis with MuECs as the sending cells and MuSCs as the receiving cells. Gene ontology (GO)-term-mediated interactome analysis identified Notch-signaling-mediated interactions between MuECs and MuSCs, including MuEC-derived Jag2 and Dll4 interacting with MuSC-derived Notch2 and Notch3(14). We verified expression of Notch signaling pathway genes via RNA-seq (Figure 1A). Freshly isolated MuECs express the Notch ligands *Dll1*, *Dll4*, *Jag1*, and *Jag2*. Freshly isolated MuSCs express the Notch receptors *Notch1*, *Notch2*, and *Notch3*, as well as the Notch ligands *Dll1* and *Jag1*, with lower expression levels of *Dll4* and *Jag2*. High expression of the Notch downstream genes *Hes1*, *Hey1*, and *Heyl* indicates that Notch signaling maintains MuSCs. We confirmed *Jag2* expression in MuECs and MuSCs using a *Jag2^LoxP/LoxP^* mouse line with a *LacZ/Neo* cassette that enabled us to detect *Jag2*-expressing cells in muscle. X-gal staining of whole tibialis anterior (TA) muscle and purified MuSCs from *LacZ/Neo*-*Jag2^LoxP/LoxP^* mice revealed high levels of β-galactosidase activity in CD31(+) capillaries, but low levels in Pax7(+) and MyoD(+) MuSCs (Figure 1B). qRT-PCR showed that *Jag2* expression is low in quiescent MuSCs, is up-regulated during activation, then down-regulated during myogenic differentiation (Figure 1C). Jag2 is detected in the membranes of cultured MuECs and MuSCs (Figure 1D). We concluded that Jag2 is expressed in MuECs and variably expressed in MuSCs, depending on their stage in myogenesis.

**Figure 1.**
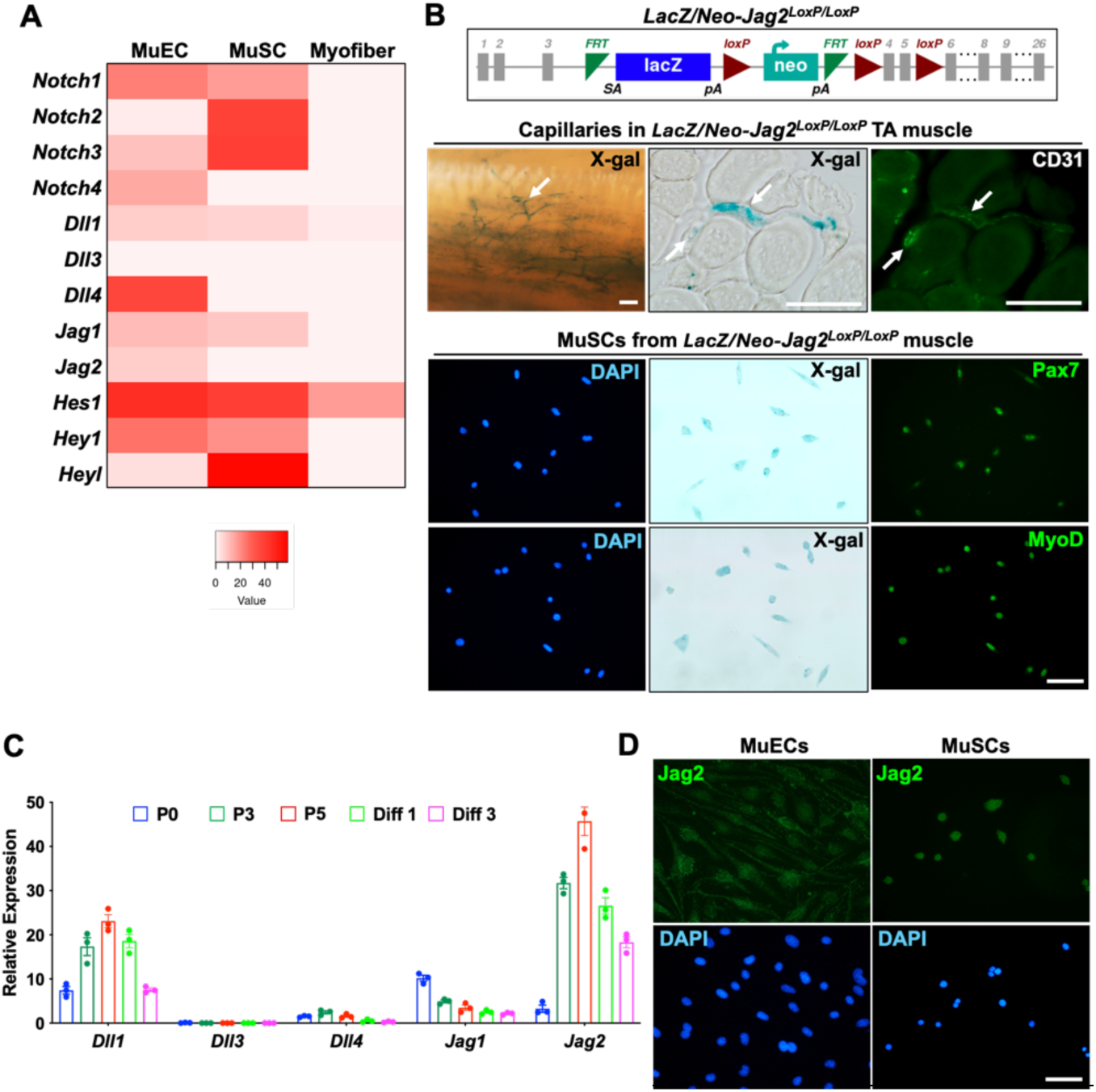
*Jag2* is expressed in MuSCs and MuECs. (A) Heatmap for gene expression by RNA-seq reveals active Notch signaling in freshly isolated MuECs and MuSCs, but not in myofibers (14). (B) Upper panel: Schematic genomic structure of *Jag2^LoxP/LoxP^* locus for conditional *Jag2* mutant (*LacZ/Neo-Jag2^LoxP/LoxP^*) mice before Flippase (Flp)-mediated recombination. Middle panels: On X-gal staining, capillaries (white arrow) in TA muscle from *LacZ/Neo-Jag2^LoxP/LoxP^* mice are positive for LacZ(+) and CD31. Lower panels: MuSCs from *LacZ/Neo-Jag2LoxP/LoxP* mice are positive for LacZ(+) and Pax7 or MyoD. (C) qRT-PCR shows upregulation and downregulation of *Dll1* and *Jag2* following MuSC activation and differentiation, respectively. P0, P3, P5, Diff 1, and Diff 3 denote freshly isolated MuSCs, passage day 3, passage day 5, differentiation day 1, and differentiation day 5. (D) Anti-Jag2 antibody staining shows Jag2 expression at the membrane of MuECs and MuSCs. DAPI stained all nuclei (blue). Scale

### Downregulation of Notch signaling and myogenic differentiation genes in *Jag2* deficiency

To understand the influence of Jag2 on Notch signaling during myogenesis, we measured the gene expression levels of 93 Notch- and myogenesis-related genes (Table S1) via qPCR in C2C12 myoblasts treated with *Jag2* shRNA compared to scrambled shRNA controls and cultured in differentiation media for 4 days. In *Jag2* shRNA treated cells, 23 genes were significantly downregulated, notably *Jag1*, *Jag2*, *Megf10*, *Notch1*, *Notch3*, and *MyoG* (Figure S1A, Table S1). No genes were significantly upregulated. After 4 days of differentiation, *Jag2* and *MyoG* expression levels were lower in *Jag2* shRNA-treated cells (Figure S1B, C), with a lower myotube fusion index (Figure S1D). Phase contrast analysis showed reduced multinucleated myotube formation in *Jag2* shRNA-treated cells (Figure S1E). These data suggest that Jag2 regulates myogenic differentiation by activating Notch signaling.

### Reduced MuSCs in homozygous *Jag2^sm^* mice

*Jag2^sm^* mice harbor a naturally occurring *p.Gly267Ser* variant in the first EGF-repeat in the extracellular domain, which is important for Notch signaling(63). Although the *Jag2^sm^*mouse displays syndactyly and cleft palate(64), a skeletal muscle phenotype has not been previously described. At 4 days after birth, the TA muscle of neonatal *Jag2^sm^* mice had small myofiber diameters (Figure 2A, B). Pax7(+) MuSCs were reduced in *Jag2^sm^* mice (Figure 2A, C). Adult *Jag2^sm^* mice showed reduced body mass and muscle weight (Figure S2). The muscle defects in neonatal *Jag2^sm^* mice persisted in adult TA muscle with reduced Feret’s fiber diameters and increased fibrosis (Figure 2A, B, D). *Jag2^sm^* mice showed reduced forelimb muscle grip strength (Figure 2E), running durations and distances (Figure 2F, G), and rotarod running time (Figure 2H). *Jag2^sm^* mice exhibit impaired MuSC development, leading to impaired skeletal muscle development and function.

**Figure 2.**
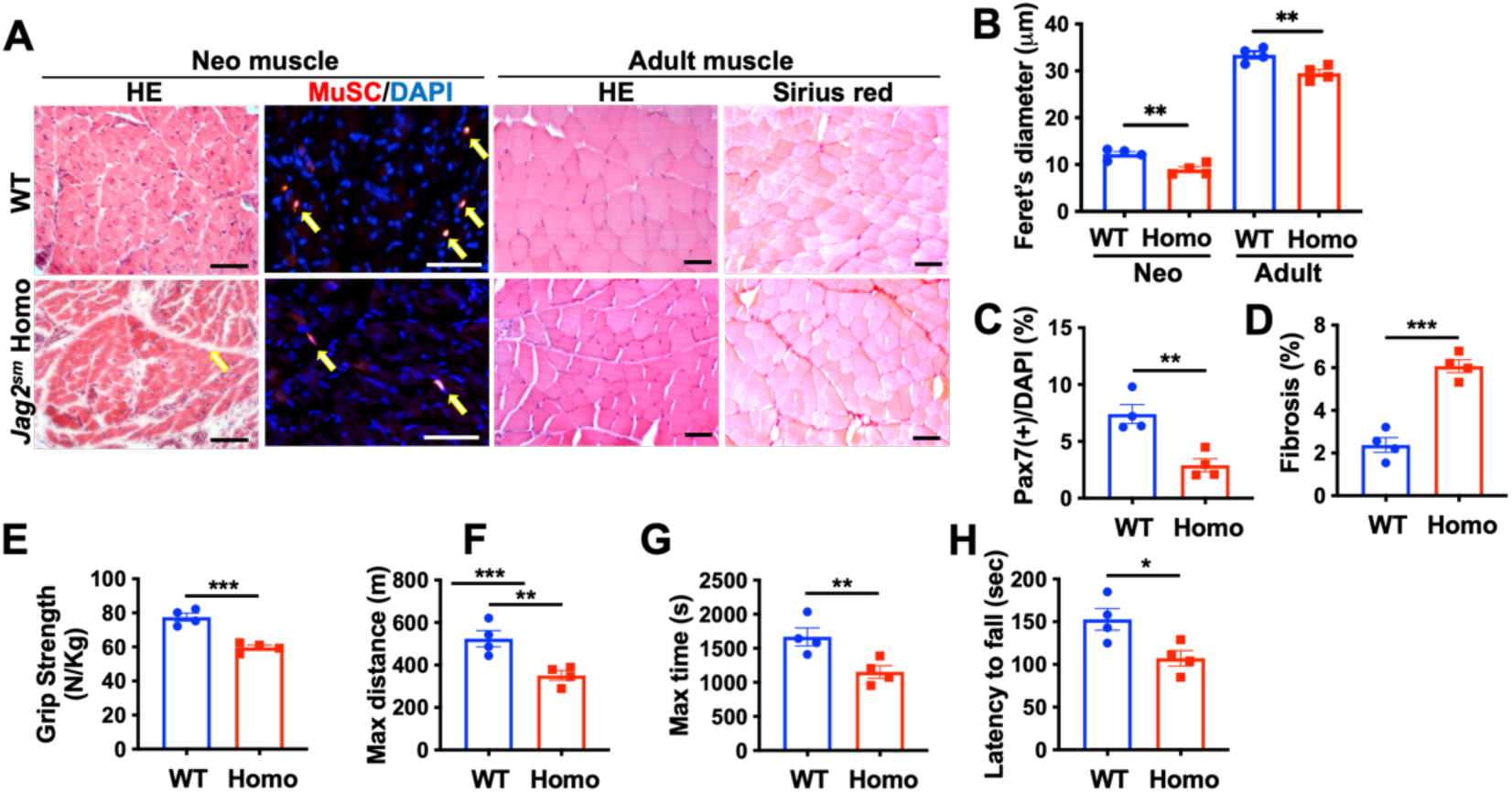
Muscle phenotypes in *Jag2^sm^* mice. (A) Muscle from 4 day-old and 3-month-old *Jag2^sm^* homozygous mice showed (B) reduced fiber diameters, (C) Pax7(+) MuSCs (arrows), and (D) increased Sirius red (+) fibrosis vs. WT mice. (E) Grip strength is reduced in *Jag2^sm^* homozygous vs. WT mice. (F, G) Treadmill running time and distance are reduced in *Jag2^sm^* homozygous vs. WT mice. (H) Motor coordination or balance on the rotarod was impaired in *Jag2^sm^* homozygous vs. WT mice. DAPI stained all nuclei (blue). Scale bars, 100*μ*m. An unpaired t-test showed *, p<0.05; **, p <0.01. ***, p <0.001. Error bars show SEM.

We crossed homozygous *Jag2^sm^* mice with *Pax7^+/CreERT2^:ROSO26^+/Loxp-stop-Loxp-tdTomato^*(*Pax7^CreERT2^:R26R^tdT^ or Pax7^tdT^*) mice to generate *Jag2^sm^ Pax7^tdT^* mice that mark MuSCs(65). We confirmed that the tdTomato was specifically expressed in the cells of interest after tamoxifen (TMX) injection (Figure 3A). Reduced number of MuSCs were detected in TA muscle cross-sections from adult *Jag2^sm^*mice (Figure 3B). Single myofiber analysis isolated from adult mice showed reduced Pax7(+) MuSC counts in *Jag2^sm^* extensor digitorum longus (EDL) (Figure 3C, D).

**Figure 3.**
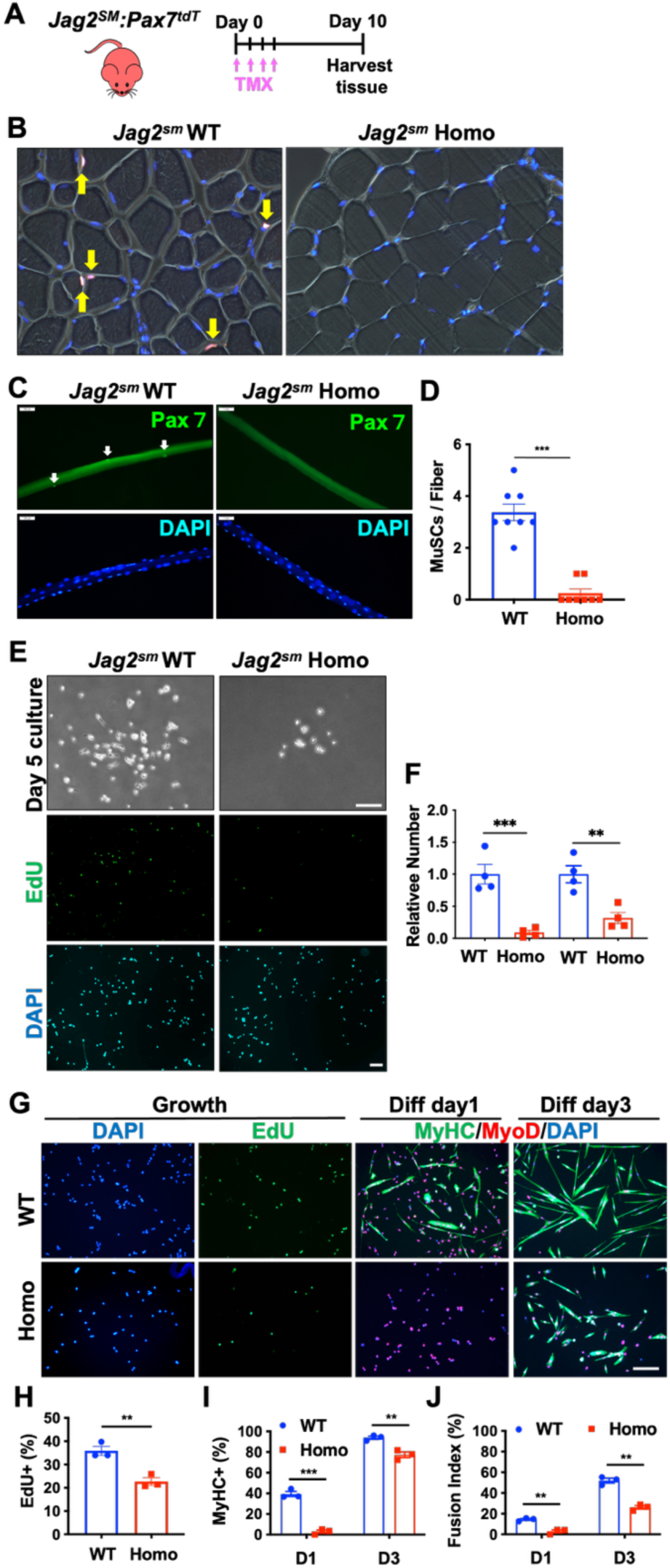
MuSCs are depleted in *Jag2^sm^*homozygous mice. (A) WT:*Pax7^tdT^*and *Jag2^sm^:Pax7^tdT^* mice were injected with TMX prior to sacrifice. (B, C, D) TA muscle sections and isolated single muscle fibers demonstrated reduced Pax7(+) MuSCs (arrows) in *Jag2^sm^* homozygous vs. WT mice. Scale bars; 20 *µ*m (top panels), 50 *µ*m (fibers). (E, F) Freshly isolated MuSCs from homozygous *Jag2^sm^* mice show reduced colony sizes and EdU(+) proliferating cells. (G) MuSCs isolated from homozygous *Jag2^sm^* mice show (H) reduced EdU(+) proliferating cells (green), (I) MyHC(+) myogenic differentiation (green) and fusion in days 1 and 3 differentiation conditions, while (J) MyoD(+) cells (red) are not altered, compared with WT cells. DAPI stained all nuclei (blue). An unpaired t-test showed **, p <0.01, ***, p<0.001. Error bars show SEM.

### Reduced cell proliferation and myogenic differentiation of MuSCs isolated from homozygous *Jag2^sm^* mice

MuSCs were isolated from hindlimb muscles of adult *Jag2^sm^*mice using antibody-mediated magnetic sorting. Freshly isolated MuSCs were cultured in growth medium to assess proliferation for 5 days. *Jag2^sm^* MuSCs displayed reduced cell proliferation, as evidenced by smaller colony sizes (Figure 3E, F). A 5-ethynyl-2’-deoxyuridine (EdU) incorporation assay correspondingly revealed reduced proliferating cells. We assessed the cell proliferation of passaged MuSCs, then switched to differentiation medium for 1 or 3 days to evaluate myogenic differentiation (Figure 3G). There were reduced proliferating [EdU(+)] MuSCs in *Jag2^sm^*(Figure 3G, H). Apoptosis of *Jag2^sm^* MuSCs was slightly increased compared with WT MuSCs following thapsigargin or UV-treatment (Figure S3). These findings confirm Jag2’s essential role in niche-independent MuSC proliferation and survival. During myogenic differentiation, the number of MyoD(+)-committed myogenic progenitors remained unchanged in homozygous *Jag2^sm^* mice (Figure 3G). However, immunostaining for myosin heavy chain (MyHC) after 1 or 3 days of differentiation showed diminished multinucleated myotubes in *Jag2^sm^*MuSC cultures (Figure 3G, I, J). These findings suggest that while *Jag2* deficiency does not affect the initial commitment of MuSCs to the myogenic lineage [MyoD(+) cells], it impairs their proliferative capacity and subsequent myogenic differentiation.

### Impaired muscle regeneration in homozygous *Jag2^sm^*mice

We assessed the regenerative capacity of TA muscles in adult *Jag2^sm^*mice following intramuscular cardiotoxin (CTX)-induced injury (Figure 4A). At 7 days post-injury, *Jag2^sm^* TA showed smaller regenerating myofibers (Figure 4B). Feret’s myofiber diameters were reduced in *Jag2^sm^*TA (Figure 4D, E) at 7- and 21-days post-injury. Immunostaining showed that embryonic myosin heavy chain (eMyHC), a marker of newly formed fibers, persisted in *Jag2^sm^*muscle at day 7 post-injury but was no longer detectable in regenerating WT muscle (Figure 4B), indicating delayed muscle regeneration in *Jag2* deficiency. Oil Red O staining revealed increased lipid accumulation in regenerating *Jag2^sm^* muscle (Figure 4F). At 28 days following sequential CTX injections in *Jag2^sm^* mice, muscle regeneration was impaired (Figure 4C, G), indicating defects in MuSC self-renewal. No differences were observed in the number of CD31(+) capillaries between *Jag2^sm^* and WT muscle (Figure 4B), indicating that the muscle regenerative defects are unlikely to be related to differences in muscle microvasculature.

**Figure 4.**
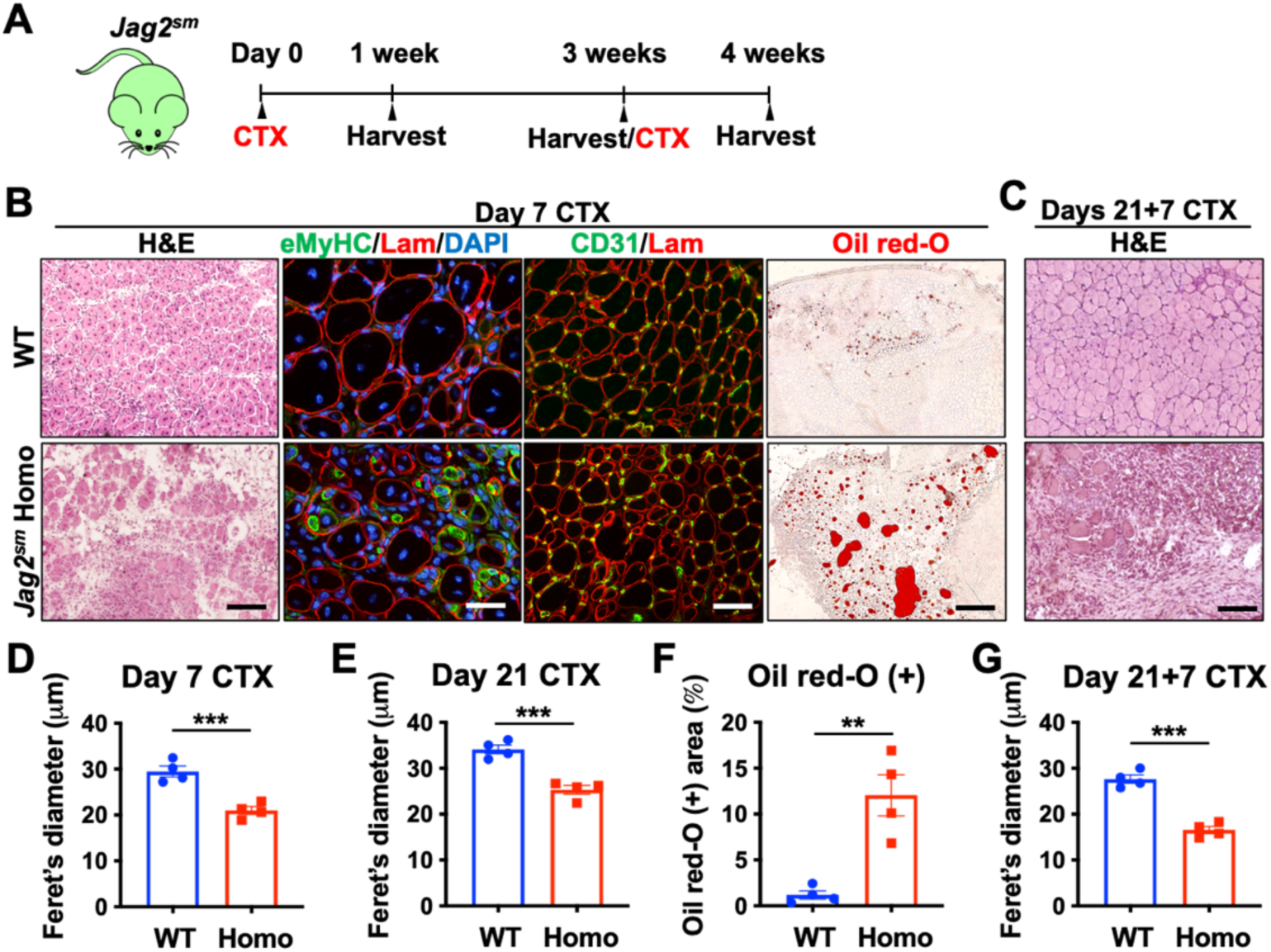
*Jag2^sm^* hypomorphic mice have muscle regenerative defects. (A) Single or repeated CTX injections into the TA muscle were performed on WT and *Jag2^sm^* mice. (B) TA histology [hematoxylin & eosin (H&E), Oil red-O] and immunostaining (eMyHC/Laminin/DAPI and CD31/Laminin) 7 days following CTX injection into the TA. Scale bars from left to right: 100, 25, 50 and 250μm. (C) H&E staining 21+7 days following sequential CTX injections into the TA. Scale bar, 100 μm. Feret’s diameters of TA fibers in WT and *Jag2^sm^* homozygous mice at (D) 7 days and (E) following CTX injection. (F) Oil red-O (+) area was evaluated at 7 days following CTX injection. (G) Feret’s diameters of TA myofibers in WT and *Jag2^sm^* homozygous mice 21+7 days following CTX injections. DAPI stained all nuclei (blue). An unpaired t-test showed **, p <0.01; ***, p<0.001. Error bars show the standard error of the mean (SEM).

### Transcriptome sequencing (RNA-seq) of homozygous *Jag2^sm^* MuSCs

To probe global gene expression changes in *Jag2* deficiency, we performed whole transcriptome sequencing (RNA-seq) on MuSCs isolated from the hindlimb muscles of *Jag2^sm^* and WT mice. MuSCs were isolated using antibody-mediated magnetic sorting. Total RNA was isolated, reverse-transcribed to cDNA, and sequenced using the Oxford Nanopore Technologies (ONT) long-read sequencing platform. Gene expression analysis revealed that 702 genes were significantly dysregulated (adjusted p-value < 0.05) in homozygous *Jag2^sm^* compared to WT MuSCs. There were 186 upregulated genes and 516 downregulated genes (Figure 5A, C, Tables S2, S3). Metascape Gene Ontology (GO) analysis indicated that 106 genes related to muscle structure development (GO:0061061), including 28 myogenic regulatory genes such as *Myog*, *Myf6*, *Mef2c*, *Mymk*, and *Igf2*, were downregulated in *Jag2^sm^* MuSCs (Figure 5B, Table S4). Negative regulators of cell proliferation, including *Cxcl12* and *Sox4*, were identified in the upregulated genes (Figure 5C, Table S2). Among Notch receptor genes, *Notch1*, *Notch2*, and *Notch3* were expressed in WT MuSCs, and *Notch2* expression was upregulated in *Jag2^sm^* MuSCs (Table S5). *Dll1*, *Jag1*and *Jag2* were detected in both WT and *Jag2^sm^* MuSCs (Table S5), suggesting *cis-*regulatory mechanisms for Notch signaling in MuSCs. Reduced myogenic differentiation and expression of Notch receptor and ligand genes was confirmed in *Jag2^sm^* MuSCs via RT-qPCR (Figure 5D). These findings are consistent with observed defects in MuSC proliferation and myogenic differentiation in *Jag2* deficiency. Increased expression of the Notch downstream effector genes *Hes1*, *Hey1*, and *Heyl*, was observed in *Jag2^sm^* compared with WT MuSCs (Figure 5D).

**Figure 5.**
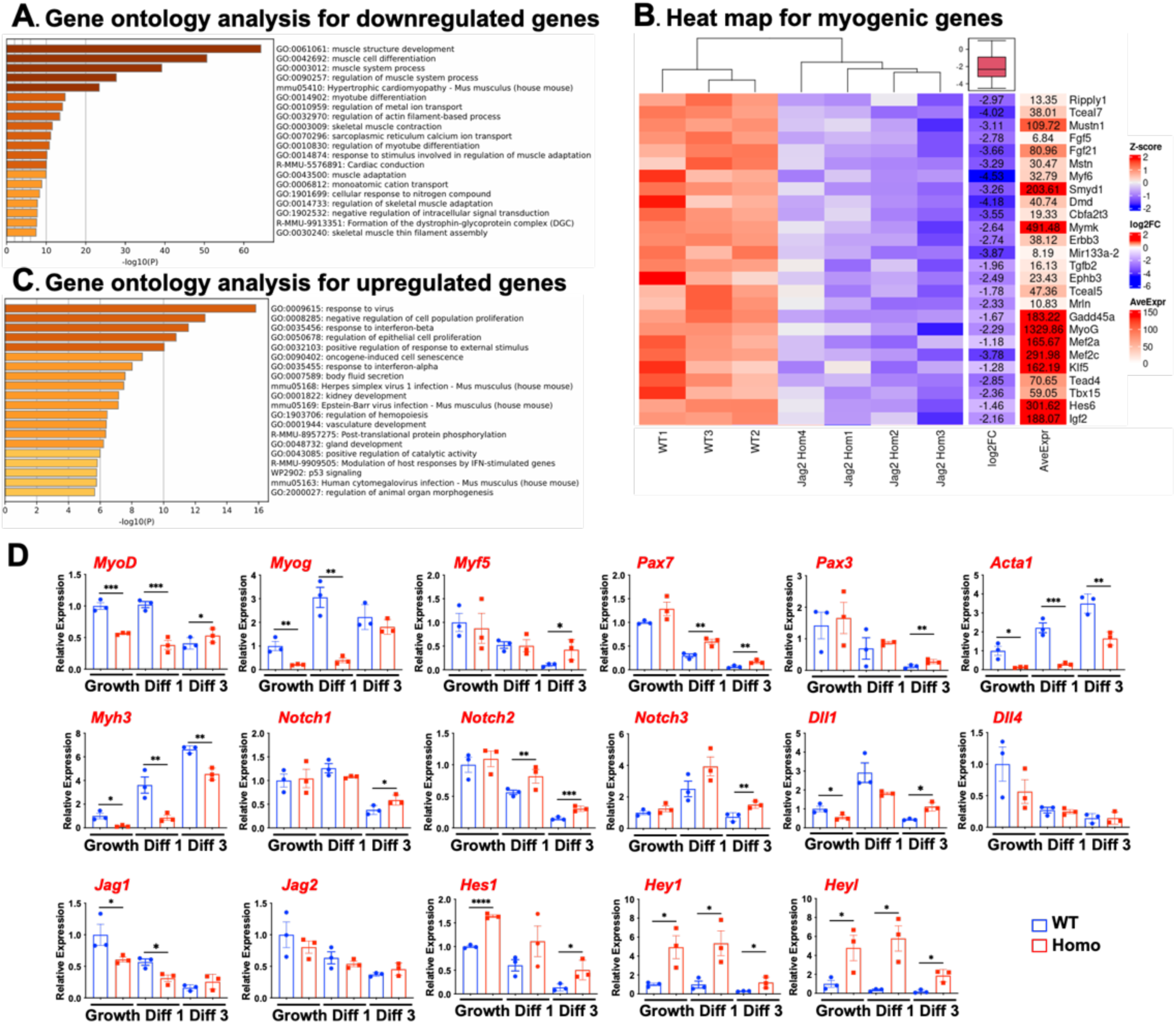
RNA-seq for gene expression profiles in *Jag2^sm^* vs. WT MuSCs. (A) Gene ontology (GO) analysis reveals that numerous significantly downregulated genes in RNA samples from *Jag2^sm^* vs. WT mouse MuSCs are muscle related. (B) Heatmap for down-regulated genes associated with myogenic regulatory genes, Notch receptor genes, and ligand genes. (C) GO analysis reveals that numerous significantly up-regulated genes in RNA samples from *Jag2^sm^*vs. WT mouse MuSCs are involved in negative regulation of cell proliferation. (D) RT-qPCR was performed on WT and *Jag2^sm^* MuSCs under growth, day 1, and day 3 differentiation conditions to detect the expression of myogenic and Notch pathway-related genes.

### *Cis-*inhibition of Notch signaling by *JAG2* in MuSCs

To determine whether human JAG2 suppresses Notch signaling in MuSCs via *cis-*inhibition, we co-transfected *JAG2* with the Notch reporter gene *pHes1-467-Luc*, containing the 467 bp *Hes1* gene upstream region, or with the *pHes1-467-RBPJ*(*-*)*-Luc*, which lacks a RBP-J binding site essential for assembly of the transcriptional complex with NICDs and other binding partners and subsequent Notch target gene activation (Figure 6A). *Hes1-467-Luc* activity was elevated in *Jag2^sm^* MuSCs, then abolished when the reporter gene with the mutant RBP-J binding site was used (Figure 6B). Notch reporter activation was blunted by N-[N-(3,5-Difluorophenacetyl)-L-alanyl]-S-phenylglycine t-butyl ester (DAPT), a global *γ*-secretase/Notch inhibitor(66). These data indicate that *Jag2* deficiency promotes Notch signaling in MuSCs. To determine which Notch receptors are targeted by Jag2, the *Hes1-467-Luc* vector was co-transfected with human *JAG2* expression vector and *Notch* expression vectors. *JAG2* suppressed Notch1, Notch2, and Notch3 but not Notch4-mediated Notch reporter gene activation (Figure 6C). Since Notch1, Notch2, and Notch3 are detected in quiescent and activated MuSCs, Jag2-mediated Notch signaling in MuSCs might be mediated through these Notch receptors. Since *JAG2* suppresses Notch signaling in MuSCs while *Jag2^sm^* MuSCs show increased Notch signaling, Jag2 may suppress Notch signaling via *cis-*inhibition. WT and *Jag2^sm^* MuSCs transfected with reference *JAG2* but not variant *JAG2* (*p.Glu164Lys*, *p.Pro682Ser*, or *p.Phe977Ser*) show reduced Notch signaling inhibition, confirming that *JAG2* pathogenic variants lack *cis-*inhibitory effects in MuSCs (Figure 6D).

**Figure 6.**
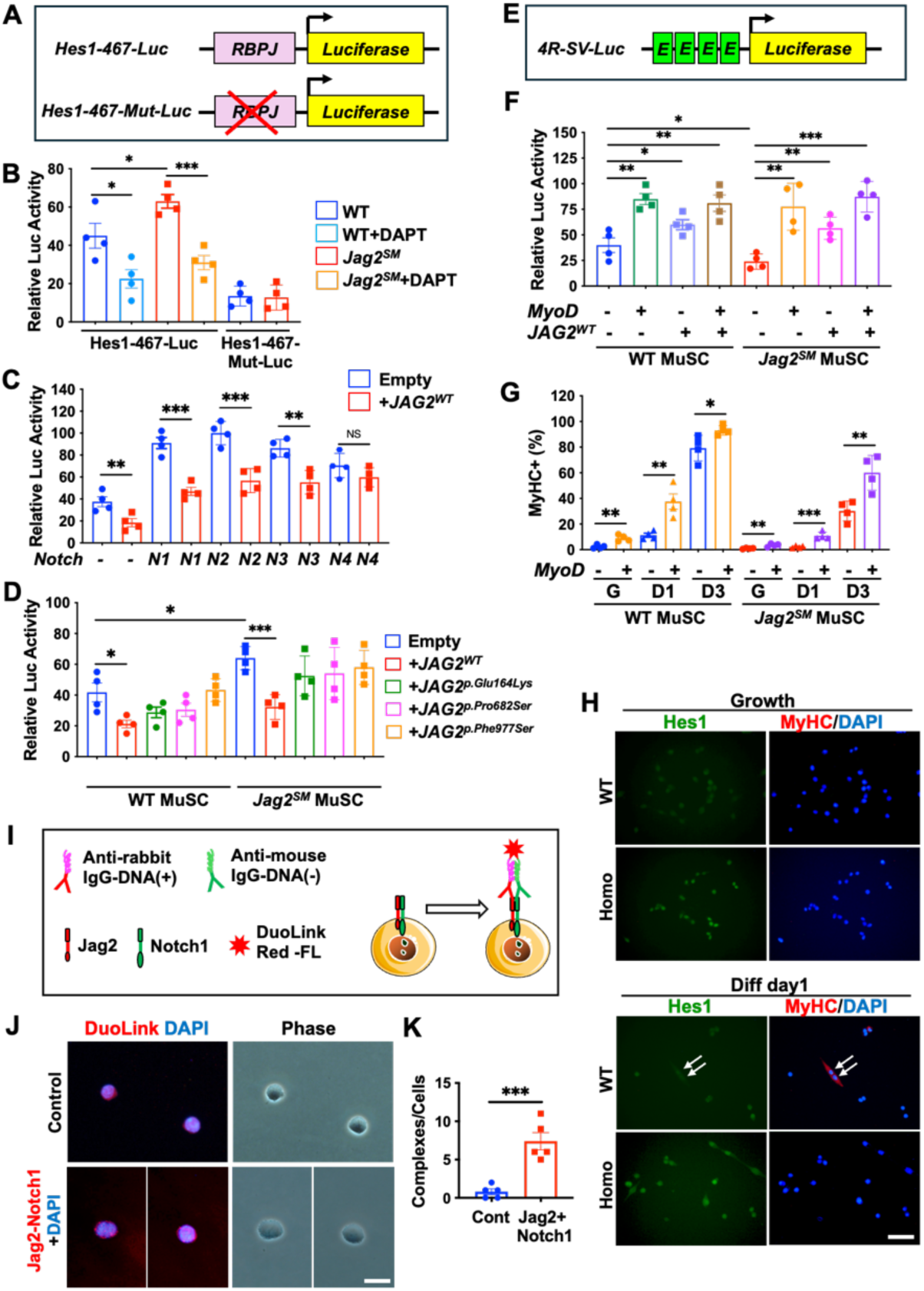
Human JAG2 suppresses Notch signaling and promotes myogenesis. (A) *Hes1-467*-Luc vector contains *Hes1* 467bp upstream promoter-driving *luciferase* (*Luc*) gene with an RBPJ binding site and *Hes1-467-Mut-Luc* vector contains a mutated RBPJ binding site. (B) Homozygous *Jag2^sm^* MuSCs show higher *Hes1-467-Luc* activity compared with WT MuSCs. Luc activities were diminished when the RBPJ-binding site was mutated or when treated with the Notch inhibitor DAPT. (C) Expression of *Notch1-4* (*N1-4*) increased *Hes1-467-Luc* activities that were suppressed by co-transfection of human *JAG2* in WT and homozygous *Jag2^sm^* MuSCs. (D) Human *JAG2-*mediated suppression was not observed when transfected with expression vectors carrying pathogenic *JAG2* variants. (E) *4R-SV-Luc* contains 4 x E-boxes for consensus binding sites for MyoD. (F) Expression of *MyoD* and human *JAG2* activates *4R-SV-Luc* in WT and homozygous *Jag2^sm^* MuSCs*. MyoD* promoted myosin heavy chain (MyHC)(+) myogenic differentiation in WT and homozygous *Jag2^sm^* MuSCs in growth (G) and differentiation conditions (days 1 and 3). (H) Anti-Hes1 antibody staining shows Hes1 is higher in homozygous *Jag2^sm^* compared with WT MuSCs. Hes1 expression is down-regulated in MyHC(+) myocytes (arrows). Scale bars, 50μm. (I) The diagram shows a DuoLink PLA assay to examine a protein complex of Jag2 and Notch1 within MuSCs. Anti-Jag2 and anti-Notch1 antibodies were used followed by anti-rabbit and anti-mouse IgG with (+) and (–) strands of oligo DNAs. Red fluorescence tags were incorporated with successful ligation. (J) DuoLink labeling shows patchy red complexes around the cell membrane regions, with no positive complexes when antibodies were eliminated as a control. (K) Quantification of DuoLink(+) complexes/cell was performed. DAPI stained all nuclei (blue). Scale bars, 100μm. An unpaired t-test showed *, p <0.05; **, p<0.01; ***, p<0.001. Error bars show the standard error of the mean (SEM).

*Jag2^sm^* MuSCs display reduced myogenic differentiation but increased Notch activity, and consequently, reduced muscle regeneration after injury. To determine whether MyoD activity is down-regulated in *Jag2^sm^* MuSCs, we tested the luciferase activity of the MyoD-binding site-driven *4R-SV-Luc* reporter gene. The *4R-SV-Luc* incorporates 4 x E-box elements and MyoD-binding motifs, which are sourced from the enhancer region of the muscle creatine kinase (*MCK*) gene (Figure 6E)(67, 68). MyoD activity is lower in *Jag2^sm^* MuSCs (Figure 6F). Co-transfection with MyoD promotes luciferase activity in *Jag2^sm^* MuSCs. *JAG2* promoted luciferase activities in both WT and *Jag2^sm^* MuSCs. *MyoD* overexpression rescued myogenic differentiation of *Jag2^sm^*MuSCs, as indicated by an increase in MyHC(+) myocytes (Figures 6G, S4). Increased Hes1 protein was detected in *Jag2^sm^* compared with WT MuSCs (Figure 6H). The Hes1 expression was abolished in MyHC(+) myocytes in WT MuSCs. These results indicate that Jag2 is a myogenic promoter that acts as a *cis-*inhibitory factor for Notch signaling.

Using DuoLink technology, an *in situ* proximity ligation assay (PLA) that identifies two proteins in close proximity, we demonstrated the presence of a Jag2-Notch1 complex on the MuSC plasma membrane (Figure 6I, J). The quantity of DuoLink reaction products was measured, confirming the existence of Jag2-Notch1 complexes exclusively when specific antibodies were used (Figure 6K). These findings suggest that Jag2 is responsible for *cis-*inhibition of MuSC Notch signaling.

### *In vitro* rescue effects of human reference *JAG2* versus variant *JAG2* in MuSCs

Following culturing in differentiation media for 4 days, overexpression of human reference *JAG2* rescued myogenic differentiation defects in *Jag2^sm^*MuSCs, while overexpression of three human *JAG2* pathogenic variants (*p.Glu164Lys*, *p.Pro682Ser*, and *p.Phe977Ser*)(30) did not (Figures 7A, B, C, S5). Similarly, overexpression of reference *JAG2* but not variant *JAG2* (*p.Glu164Lys*, *p.Pro682Ser*, or *p.Phe977Ser*) rescued myogenic cell fusion in *Jag2*-deficient C2C12 myoblasts (Figure S6). These results demonstrate the essential roles of *JAG2* in MuSC function, providing evidence for a loss of function (LOF) pathogenic mechanism of *JAG2* variants.

**Figure 7.**
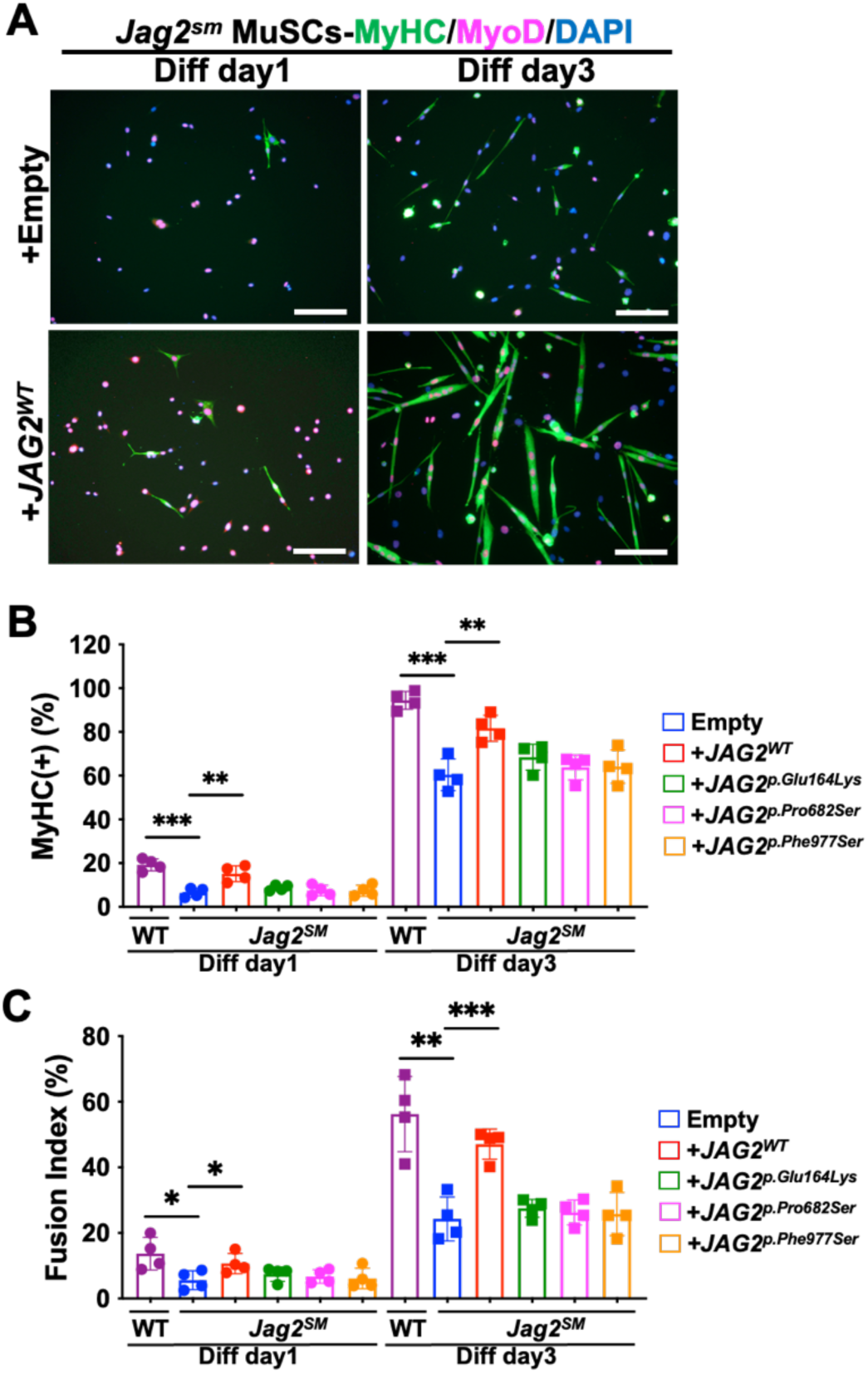
Overexpression of human *JAG2* rescues differentiation defects in *Jag2^sm^* MuSCs. MuSCs isolated from homozygous *Jag2* mice were used for expression vector-mediated human *JAG2* overexpression. (A) Overexpression of WT human *JAG2* (*JAG2^WT^*) but not human *JAG2* pathogenic variants (*p.Glu164Lys*, *p.Pro682Ser*, and *p.Phe977Ser*) increased MyHC(+) myogenic differentiation (green) (B), and fusion in days 1 and 3 differentiation conditions (C), compared with control empty vector-transfected cells. DAPI stained all nuclei (blue). Scale bars, 100 μm. An unpaired t-test showed *, p <0.05; **, p<0.01. Error bars show the standard error of the mean (SEM).

### Jag2-mediated regulation of MuSC self-renewal via MuECs

MuECs play an essential role in MuSC self-renewal during muscle regeneration (14), but the exact signaling mechanism between the two cell types is unclear. We examined Jag2-mediated effects of MuECs on MuSC self-renewal. We confirmed *Jag2* expression in MuECs and MuSCs via RNA-seq (Figure 1A), histology using a *Jag2^LoxP/LoxP^* mouse line with a *LacZ/Neo* cassette that enabled us to detect *Jag2*-expressing cells (Figure 1B), RT-qPCR (Figure 1C) and immunostaining (Figure 1D) in muscle and dissociated cells. We performed co-culture experiments using MuECs and MuSCs isolated from WT mice. MuECs were transfected with either *Jag2* or scrambled control siRNA, and their ability to support MuSC self-renewal was evaluated by quantifying Pax7(+)/MyoD(-) cells (i.e., self-renewing MuSCs or reserve cells) after 5 days of co-culture (Figure 8A). MuSCs co-cultured with *Jag2*-depleted MuECs exhibited reduced Pax7(+)/MyoD(-) self-renewing reserve cells compared to those co-cultured with control MuECs (Figure 8B, C, D). This effect was replicated in co-cultures treated with DAPT, suggesting that Jag2-mediated Notch activation in MuSCs is required for self-renewal (Figure 8C, D). These results demonstrate that Jag2 in MuECs regulates MuSC self-renewal through *trans-*activation signaling, in addition to the cell-autonomous *cis-*inhibition effects of Jag2 (Table 1).

**Figure 8.**
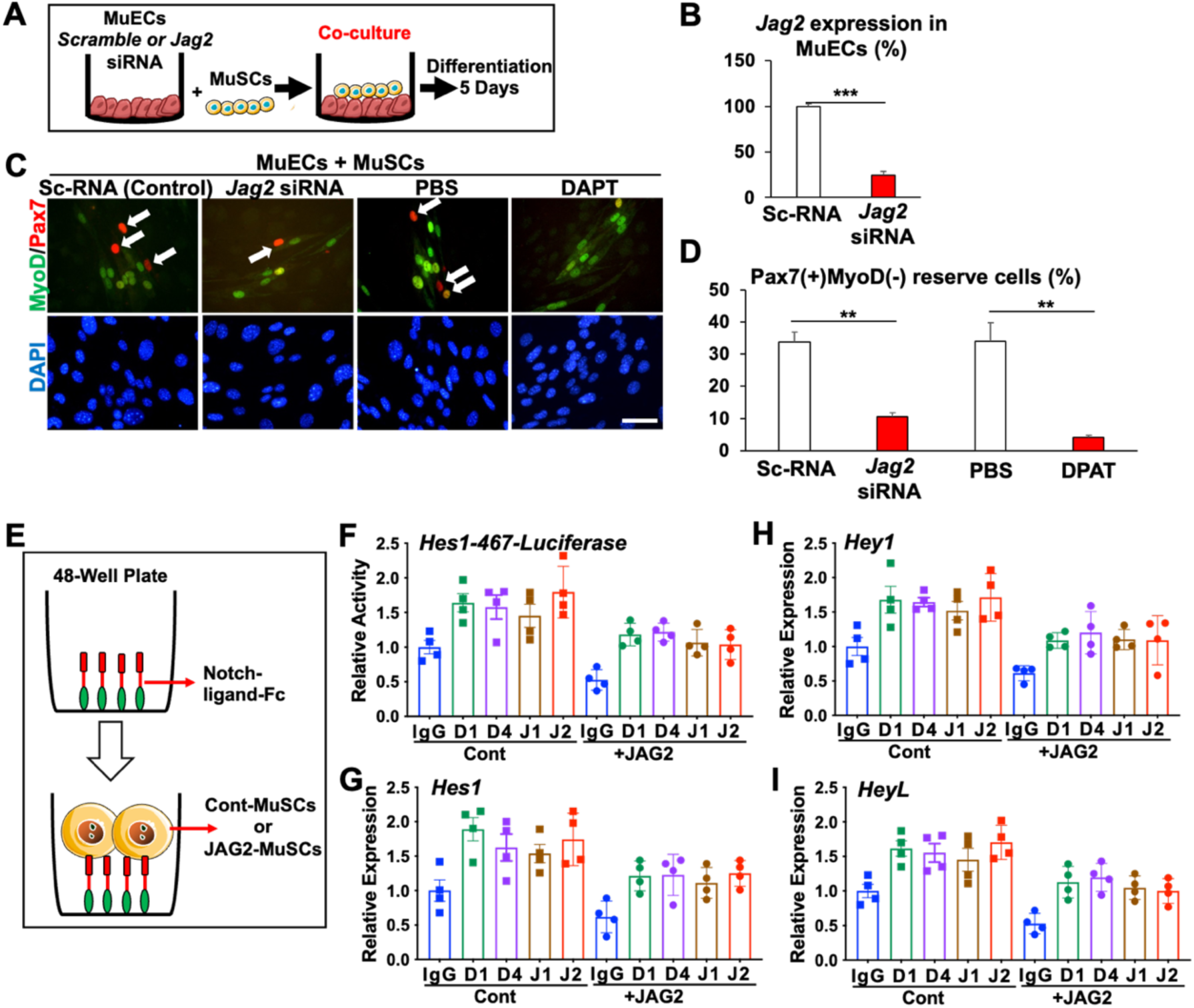
MuSC and MuEC co-culture experiments. (A) MuSCs were layered on top of the MuECs with scrambled or *Jag2* siRNA, allowed to adhere, and then co-cultured in differentiation medium for 5 days. (B) MuECs transfected with *Jag2* siRNA show a significant reduction of *Jag2* expression vs. scrambled siRNA. (C, D) Pax7(+)MyoD(-) self-renewing MuSCs were reduced when *Jag2* was co-cultured with Jag2-KD MuECs vs. control MuECs (arrows). Downregulation of Notch signaling through the pan-Notch inhibitor DAPT reduced the number of Pax7(+)MyoD(-) self-renewing MuSCs vs. PBS-treated cells in the co-cultures (arrows). (E) Diagram of the evaluation of MuSCs treated with Notch ligands. (F) Hes1-467-Luciferase activity was assessed in control and *JAG2*-expressing MuSCs exposed to Notch ligand (Control-IgG-Fc, Dll1-Fc, Dll4-Fc, Jag1-Fc, and Jag2-Fc). Comparative mRNA expression levels of the Notch effector genes (G) *Hes1*, (H) *Hey1*, and (I) *HeyL*, in control and *JAG2*-expressing MuSCs exposed to Notch ligand (Control-IgG-Fc, Dll1-Fc, Dll4-Fc, Jag1-Fc and Jag2-Fc). DAPI stained all nuclei (blue). An unpaired t-test showed **, p<0.01; and ***, p<0.001. Error bars show the standard error of the mean (SEM). Scale bars, 50 μm.

**Table 1.** Muscle phenotypes for *Jag2* deficiency in various settings.

| Cell or tissue type model<br>*Jag2* effect | MuSCs (mouse)<br>or<br>AMPs (fly) | MuECs<br>(mouse) | Skeletal muscle<br>(mouse) |
| --- | --- | --- | --- |
| *Jag2<sup>sm</sup>* mouse<br>(hypomorphic)<br>*Cis-inhibition*<br>*Trans-activation* | Reduced number of<br>MuSCs<br>Reduced<br>proliferation and<br>differentiation | No observed<br>phenotype in<br>capillaries | Reduced fiber<br>diameter and<br>increased fibrosis<br>Impaired muscle<br>regeneration<br>following muscle<br>injury |
| Mouse with MuSC-specific<br>*Jag2* KO<br>(*Jag2<sup>LoxP/LoxP</sup>:Pax7<sup>CreERT2</sup>*)<br>*Cis-inhibition* | Reduced<br>proliferation and<br>differentiation<br>Normal MuSC self-renewal | N/A | Reduced fiber<br>diameter Reduced<br>muscle regeneration<br>following muscle<br>injury |
| Mouse with MuEC-specific<br>*Jag2* KO<br>(*Jag2<sup>LoxP/LoxP</sup>:VEcad<sup>CreERT2</sup>*)<br>*Trans-activation* | Reduced MuSC self-renewal following<br>sequential muscle<br>injuries | N/A | Reduced fiber<br>diameter Impaired<br>muscle regeneration<br>following sequential<br>muscle injuries |
| siRNA-mediated *Jag2* KD<br>in MuECs<br>(co-culture with MuSCs)<br>*Trans-activation* | Reduced MuSC self-renewal | N/A | N/A |
| *Serrate* RNAi flies<br>*Trans-activation* | Decreased AMP<br>proliferation(69) | N/A | Decreased<br>locomotor activity<br>(including flight and<br>gait) in adult flies |

To determine whether extracellular Jag2 can *trans-*activate endogenous Notch activity and whether that activity is suppressed by intracellular Jag2 via *cis-*inhibition, WT MuSCs were plated onto dishes that were treated with Notch ligands linked to the Fc domain of IgG: Dll1-Fc, Dll4-Fc, Jag1-Fc, and Jag2-Fc (Figure 8E). Anti-goat IgG was a control. Dll1-Fc, Dll4-Fc, Jag1-Fc and Jag2-Fc increased *Hes1-467-Luciferase* activities, which were suppressed by *JAG2* expression in WT MuSCs (Figure 8F), prompting elevated expression of the Notch effector genes *Hes1*, *Hey1* and *HeyL*, compared with control-Fc treatment (Figure 8G-I). These extracellular Notch-ligands mediated the up-regulation of *Hes1*, *Hey1*, and *HeyL*, which was suppressed by *JAG2*. Therefore, Dll1, Dll4, Jag1, and Jag2 *trans-*activate Notch signaling and promote self-renewal in MuSC cultures, which are suppressed by intracellular Jag2-mediated *cis-*inhibition.

### MuECs-derived *Jag2* is essential for MuSC self-renewal, while MuSC-derived *Jag2* is essential for proper MuSC myogenic differentiation

Our RNA-seq analysis and *in vitro* experiments demonstrated that Jag2 regulates MuSC proliferation, myogenic differentiation, and self-renewal via both cell-autonomous *cis-*activation and MuEC-MuSC-interaction-mediated *trans-*activation. To confirm these findings *in vivo*, we investigated the effects of MuEC- and MuSC-specific *Jag2* deletions mediated by *VE-cadherin^CreERT2^* and *Pax7^CreERT2^*, respectively, using the *VEcad^CreERT2^:Jag2^LoxP/LoxP^:Pax7^tdT^*and *Pax7^CreERT2^:Jag2^LoxP/LoxP^:Pax7^tdT^* mice following TMX-injection (Figures 1B, 9A). TA muscles were harvested 7 days and 7+21days following CTX injections. Using qPCR, we verified efficient conditional *Jag2* gene deletion (*Jag2^Δ/Δ^*) in MuECs and MuSCs following Cre activation, in *VEcad^CreERT2^:Jag2^LoxP/LoxP^:Pax7^tdT^*and *Pax7^CreERT2^:Jag2^LoxP/LoxP^:Pax7^tdT^* mice, respectively (compared to control *Jag2^+/+^* MuECs and MuSCs) (Figure S7). TA cross-sections in VEcad-mediated *Jag2* conditional knockout (cKO) mice (*MuEC-Jag2^Δ/Δ^*) displayed reduced muscle regeneration 7 days following CTX injection (Figure 9B). At 28 days following sequential CTX injections in *MuEC-Jag2^Δ/Δ^* mice, muscle regeneration was impaired (Figure 9B, C), indicating defects in MuSC self-renewal. MuSCs were reduced in the regenerating TA at 3 weeks (Figure 9E, F), confirming that self-renewal of MuSCs is regulated by MuEC-derived Jag2 during muscle regeneration. The reduced MuSCs in fully repaired muscle suggest a failure of MuSC self-renewal in *MuEC-Jag2^Δ/Δ^*mice. MuSC self-renewal was reduced in MuEC-specific *Jag2* knockout mice, indicating that Jag2 is essential for MuSC self-renewal via MuEC-mediated *trans-*effects during muscle regeneration (Table 1).

**Figure 9.**
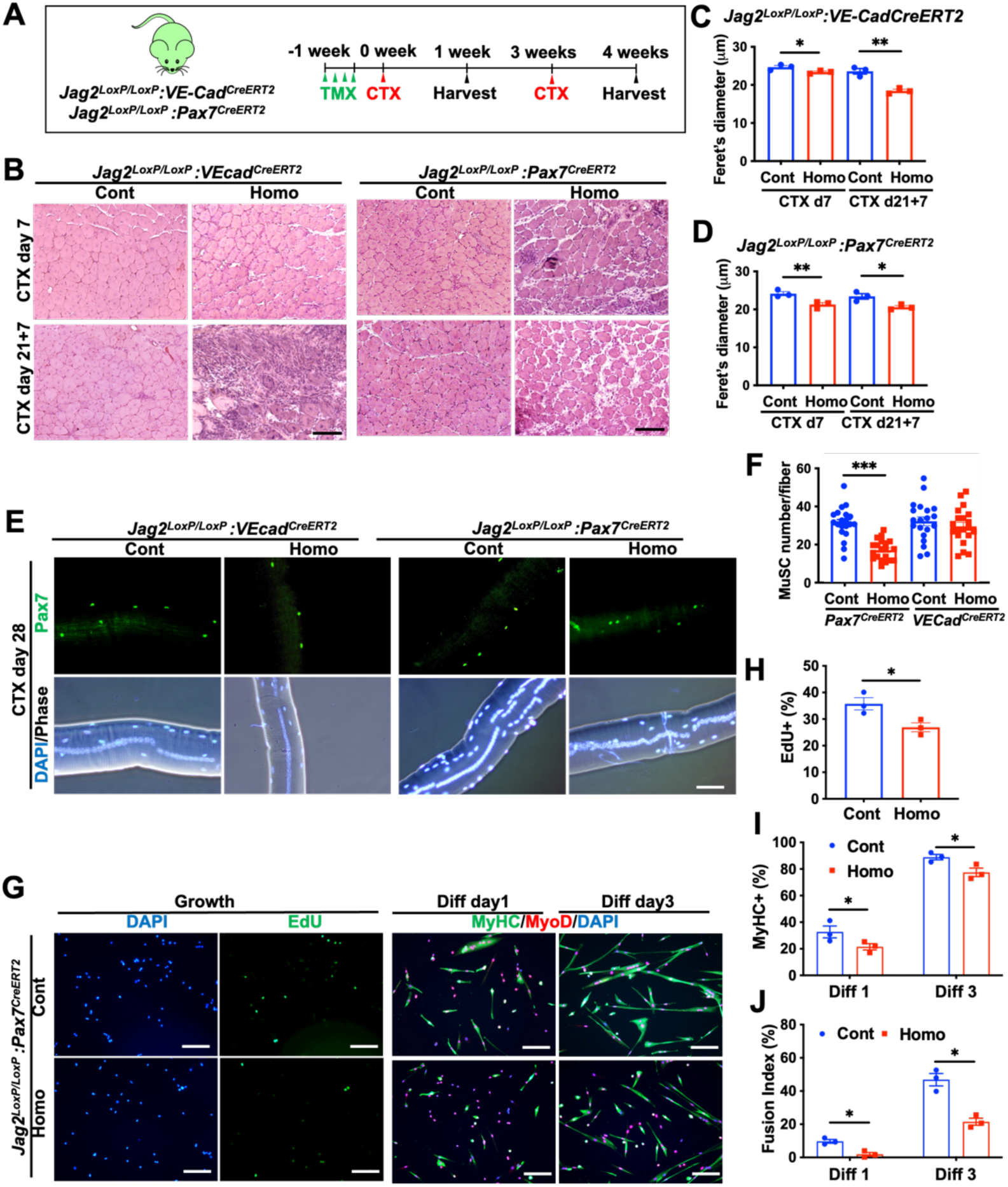
Reduced self-renewal in *Jag2^LoxP/LoxP^:VEcad^CreERT2^* and reduced regeneration in *Jag2^LoxP/LoxP^:Pax7* mice. (A) Following tamoxifen (TMX) injection, single or repeated CTX injections into the TA were performed. (B) H&E staining of TA by 7 days following CTX injection or 28 days following sequential CTX injections in *Jag2^LoxP/LoxP^:VEcad^CreERT2^* and *Jag2^LoxP/LoxP^:Pax7^CreERT2^* mice. Scale bars, 100 μm. (C, D) Feret’s diameters of TA muscle fibers in *Jag2^LoxP/LoxP^:VEcad^CreERT2^* and *Jag2^LoxP/LoxP^:Pax7^CreERT2^* mice following CTX injections. (E, F) Single muscle fibers were isolated at 28 days following TMX treatment and CTX injection. Anti-Pax7 antibody staining shows a reduced number of self-renewing MuSCs in *Jag2^Δ/Δ^:VEcad^CreERT2^* but not in *Jag2^Δ/Δ^:Pax7^CreERT2^* mice. (G) MuSCs isolated from homozygous *Jag2^Δ/Δ^:Pax7^CreERT2^* mice with TMX treatment show (G, H) reduced EdU(+) proliferating cells (green) in growth, (G, I) MyHC(+) myogenic differentiation (green), and (G, J) fusion index in day 1 and 3 of differentiation conditions compared with control cells. DAPI stained all nuclei (blue). Scale bars, 50 μm for (E) and 100 μm for (G). An unpaired t-test showed *, p<0.05; **, p<0.01; and ***, p<0.001. Error bars show the standard error of the mean (SEM).

TA cross-sections in Pax7-mediated *Jag2* cKO (*MuSC-Jag2^Δ/Δ^*) mice displayed reduced muscle regeneration by 1, 2, and 3 weeks following single CTX injections (Figure 9B, D). However, MuSC numbers were not altered in regenerating TA by 3 weeks (Figure 9E, F), indicating that self-renewal of MuSCs is not regulated by MuSC-derived Jag2 (i.e., *cis-*activation) during muscle regeneration. Isolated single muscle fibers from *MuSC-Jag2^Δ/Δ^* mice also showed similar numbers of MuSCs compared with those in *MuSC-Jag2^+/+^*mice (Figure 9E, F). MuSCs cultured from *Jag2* MuSC-specific knockout mice show reduced proliferation and differentiation (Figure 9G, H-J). We conclude that Jag2 is essential for MuSC differentiation via cell-autonomous *cis-*inhibition of Notch signaling during muscle regeneration (Table 1).

### Human *JAG2* rescues *Serrate* deficiency in *Drosophila*, but pathogenic variants do not

*Serrate* (*Ser*), the *Drosophila* ortholog of human *JAG1* and *JAG2*, plays a critical role in wing development. In the wing disc, *Serrate* is expressed in the epithelium adjacent to adult muscle progenitor cells (AMPs), where it may promote the proliferation of muscle progenitors via *trans-* activation of Notch(69). Ser is also expressed in a distinct subset of AMPs(70). We generated transgenic fly lines carrying UAS-human *JAG2* constructs, allowing for tissue-specific expression of *JAG2* using the Gal4/UAS system. When reference human *JAG2* (*JAG2^Ref^*) was overexpressed using the *Serrate-Gal4* driver mimicking the endogenous expression pattern of *Serrate*, flies developed normally. However, the wings in the corresponding adult transgenic flies showed a characteristic "delta" wing vein phenotype(71), indicating a genetic interaction between human *JAG2* and the *Drosophila* Delta-Notch pathway (Figure 10A). Expression of either human pathogenic *JAG2* variant (*p.Glu164Lys* or *p.Pro682Ser*) associated with muscular dystrophy in humans(30) led to an attenuated wing vein delta phenotype, suggesting a LOF effect (Figure 10A).

**Figure 10.**
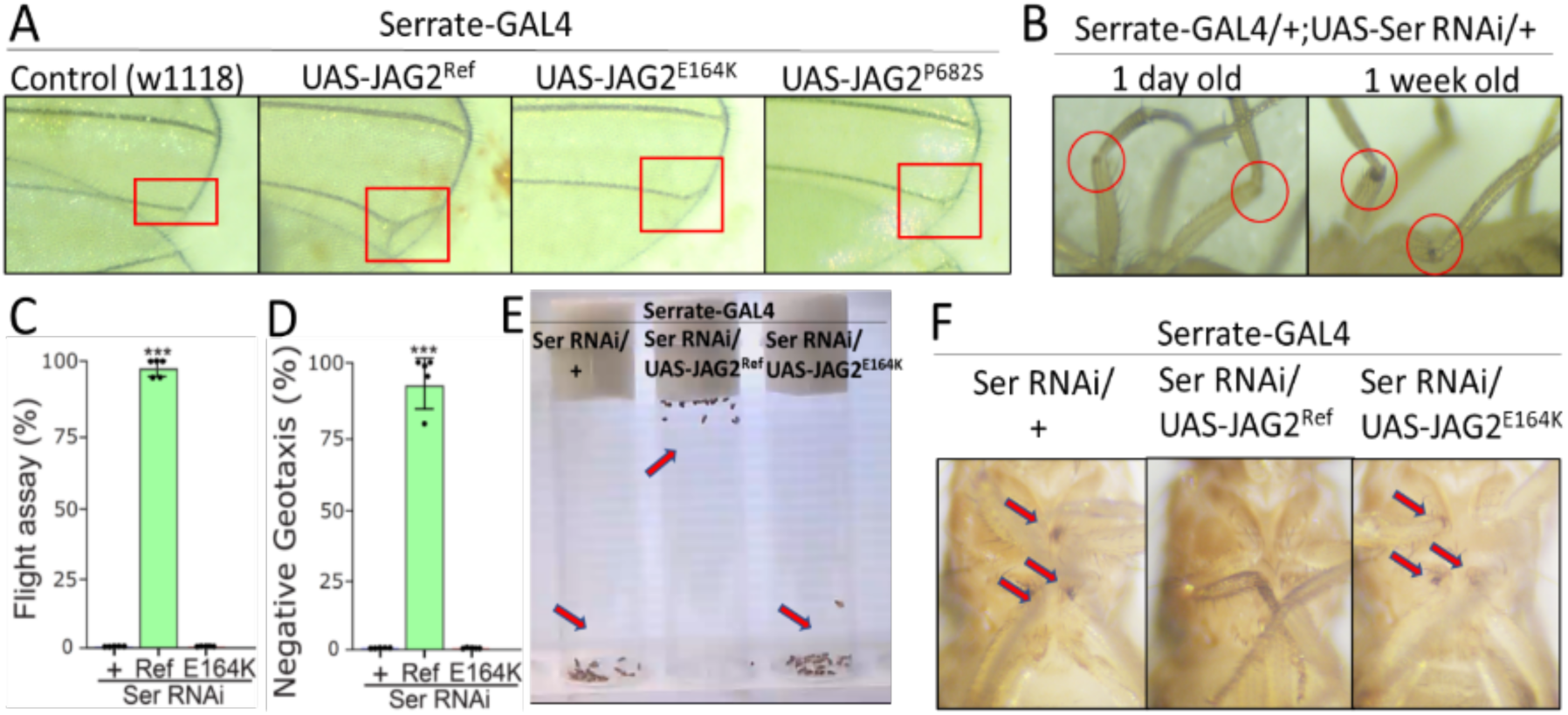
Reference human *JAG2* (*JAG2^Ref^*) rescues Serrate deficiency in *Drosophila*, while human variant *JAG2^E164K^* does not. (A) Human *JAG2^Ref^* induced delta-shaped wing veins,but variants showed marginal effects. (B) Serrate deficiency generated progressive melanotic spots on the legs. Expression of *JAG2^Ref^* rescued manifestations of serrate deficiency, whereas expression of *JAG2^E164K^* did not, on measures of (C) flight and (D, E) negative geotaxis, along with (F) melanotic spots. n=5 replicates, ***, p<0.001 (two-sided t-test).

Likewise, our results in mice showed that the pathogenic *JAG2* variants result in LOF (Figures 2-4, 9, and Table 1). RNA interference (RNAi) suppressed endogenous *Serrate* expression in *Serrate*-expressing cells with *Serrate-Gal4*. RNAi-mediated Ser knockdown did not induce developmental abnormalities, such as pupal lethality or eclosion defects. Right after eclosion, the Ser RNAi adult flies displayed normal walking and flight behavior. However, a rapid decline in locomotor activity occurred within a week post-eclosion, including both flight and gait impairments (Figure 10B-E). Flies with reduced *Serrate* exhibited progressive development of dark melanotic spots, indicating tissue necrosis and a hemocyte-mediated inflammatory response. This is consistent with the need for Notch signaling in the leg imaginal discs to promote leg segment formation(72). The degenerative phenotypes observed with *Serrate* LOF were useful to assess whether human *JAG2* could rescue the defects. Expression of reference human *JAG2*, but not *JAG2 p.Glu164Lys*, in *Serrate*-deficient flies rescued the locomotor deficits and necrotic legs (Figure 10C-F). These findings demonstrate the functional conservation of *JAG2* across species and provide *in vivo* evidence for the LOF mechanism of pathogenic *JAG2* variants (Table 1).

## DISCUSSION

Biallelic pathogenic variants in the canonical Notch ligand *JAG2* cause a form of muscular dystrophy(30, 73, 74). Pathogenic variants in the paralogous gene *JAG1* are associated with Alagille syndrome(75, 76), which does not prominently involve skeletal muscles, yet *JAG1* augmentation shows promise as a therapeutic target for muscular dystrophy(77, 78). Notch signaling affects several biological functions associated with skeletal muscle, including MuSC self-renewal, maintenance, and muscle regeneration. Jag2 is expressed in both MuSCs and MuECs, but the Significance of Jag2 for skeletal muscle development and health was not clear.

Our data indicate that *Jag2* deficiency in MuSCs impairs their myogenic differentiation potential via failed *cis-*inhibition effects for Notch activity, while *Jag2* deficiency in MuECs impairs MuSC self-renewal via failed *trans-*activation effects for Notch activity as niche cells, suggesting that Jag2-related cell-autonomous (*cis*) and cell-nonautonomous (*trans*) Notch signaling affects skeletal muscle development, regeneration and health in different ways (Figure 11). In *Drosophila*, *Serrate*-expressing niche cells in the wing epithelium regulate the proliferation and maintenance of adult muscle precursors (AMPs)(69). We demonstrated that *in vivo* knockdown of *Serrate* (the fly ortholog of *JAG2*) in *Serrate*-expressing cells resulted in motor function and morphological defects. These phenotypes are postulated to result from reduced *trans-* activation. It is unclear if *cis-*inhibitory activity is also involved in Drosophila adult muscle development (Figure 11).

**Figure 11.**
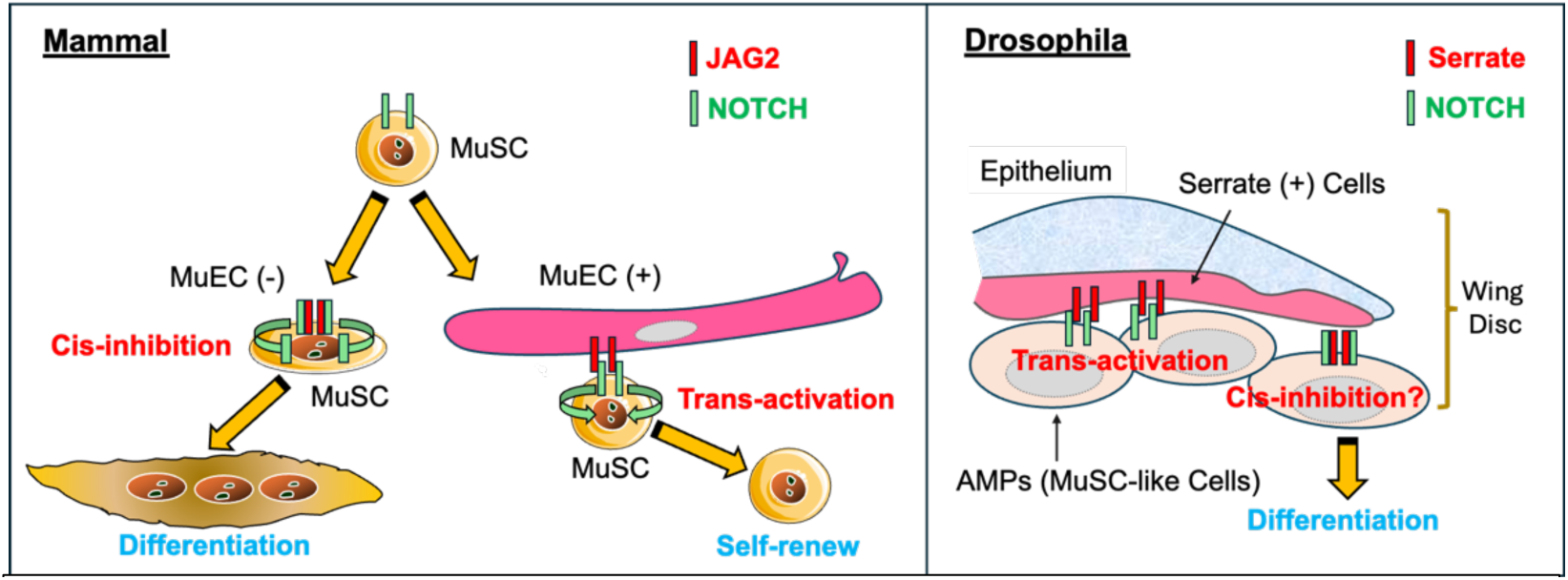
A diagram illustrating the role of *Jag2* expression in mammal (left) and *Drosophila* (right) muscles. In mammals, neighboring capillary MuECs *trans*-activate Notch signaling in MuSCs via *Jag2* for MuSC self-renewal. MuSCs, which do not receive *Jag2*-mediated *trans*-activation by MuECs suppress Notch signaling via *cis-*inhibition by cell-autonomous *Jag2* expression, stimulating myogenic differentiation. In *Drosophila* wing discs, the ortholog *Serrate* is expressed in epithelial cells, which activates Notch signaling in adjacent adult muscle precursors (AMPs), which are MuSC-like cells, to maintain the progenitor pool. AMPs express *Serrate*, but it is unclear whether *cis-*inhibition by *Serrate* occurs in AMPs.

Homozygous *Jag2* hypomorphic (*Jag2^sm^*) mice display digit and craniofacial developmental defects(79, 80). We showed that homozygous *Jag2^sm^* mice display impaired muscle regeneration due to a reduced Pax7-positive MuSC population during muscle development. The surviving homozygous *Jag2^sm^* MuSCs showed reduced proliferation and decreased myogenic differentiation. To reveal whether MuSC defects seen in *Jag2^sm^*mice are due to the *trans-*effects via neighboring niche cells or cell-autonomous *cis-*effects, we utilized MuSC-MuEC co-culture experiments, Notch ligand-Fc treatment cultures, and Cre-recombinase-mediated conditional *Jag2* gene KO mice. We demonstrated that MuEC-specific *Jag2* knockout resulted in reduced MuSC self-renewal. Our MuEC and MuSC co-culture experiments demonstrated that MuEC-derived Jag2 is required for sufficient MuSC self-renewal, underscoring the significance of direct cellular interaction between MuECs and MuSCs for the activation of Notch signaling *in vitro* and *in vivo*.

### *Trans-*activation of Notch signaling by Jag2

Multiple investigations have implicated MuECs in the functionality of MuSCs(11–13). We and others have demonstrated that vascular network enhancement augments MuSC populations in mouse models of Duchenne muscular dystrophy (DMD)(15–17, 81). Recent works, including our studies, reveal a molecular mechanism that connects MuECs and MuSCs, along with the functional outcomes of this signaling. We demonstrated that the proximity of MuSCs to capillaries is actively orchestrated by VEGFA secreted by MuSCs, attracting capillaries to create a juxtavascular environment for MuSCs(14). In addition, MuSC self-renewal is induced by Notch activation, which is stimulated by the adjacent capillary MuECs through Dll4 as a *trans-*activator. By contrast, the MuEC-derived secreted form of Dll4 regulates muscle fiber atrophy(82). Our *in vitro* co-culture experiments and *in vivo* conditional gene KO mice showed that MuEC-derived Jag2 is essential for MuSC self-renewal. Therefore, MuEC-mediated MuSC self-renewal requires at least two Notch ligands, Dll4 and Jag2.

Recent findings underscore the necessity of Notch receptors for MuSCs to revert to quiescence and establish stem cell populations(83, 84). Regarding the neighboring cell origin for Notch activation, Dll4, derived from mature myofibers, activates Notch3 expression in MuSCs, facilitating their return to quiescence(84). Mature myofiber-derived Dll4 is important for the maintenance of quiescent MuSCs on myofibers in myofiber-specific *Dll4* KO mice(85). Moreover, Dll4 and Jag2 from muscle fibers regulate Notch signaling in the proximal MuSCs to enhance their regenerative potential via increased self-renewal(86). We found that Dll4 and Jag2 levels were reduced in myofibers compared to MuECs, suggesting that MuECs play a crucial role as Notch ligand-synthesizing cells that support MuSC self-renewal in skeletal muscle.

### *Serrate-Notch*-mediated *Drosophila* myogenic progenitor maintenance

Using *Serrate-GAL4* to knock-down *Serrate* in *Serrate*-expressing cells, we observed previously unreported adult *Drosophila* phenotypes, which we believe are due to reduced *trans-*activation. Whether Drosophila *Serrate* is also involved in *cis-*inhibition is unknown. Recent single-nucleus sequencing(87) suggests that a discrete population of *Serrate*-expressing cells is present in the adult muscle system, although these are fewer than *Delta*-expressing cells and do not co-express *Delta*, and thus remains the main *cis-*inhibitory signal in fruit flies.

### *Cis-*inhibition

We demonstrated that MuSC-specific *Jag2* knockout resulted in reduced myogenic differentiation without affecting MuSC self-renewal capacity. These results are consistent with our RNA-seq and gene knockdown data. Notch signaling relies on families of ligands and receptors that relay messages to adjacent cells in various combinations across distinct cell types, as seen in MuEC-MuSC interactions (in *trans*), and within the same MuSCs (in *cis*). Notch ligands and their corresponding receptors that are present within the same cell display *cis-* inhibition of Notch signaling(88–90). This *cis-*inhibition plays a crucial role in various developmental processes, such as wing disc formation in *Drosophila*, maintenance of epidermal stem cells, neurogenesis, pancreatic cell differentiation, and hematopoiesis (58, 88, 91). It has been proposed that Notch ligands are capable of binding to Notch receptors and of *cis-*activation of Notch signaling within the same cells(59, 92). The interaction between the Notch receptor and its ligand arises from *trans-*activation and *cis-*activation of the Notch receptor mediated by the ligand as a monomer, while *cis-*inhibition is induced by the ligand as a dimer(90). Therefore, *cis-* activation indicates innovative methods through which cells can integrate various Notch interactions. However, more recent work demonstrated that with competition from *trans-*activating ligands from neighboring cells, Dll1 and Dll4 can *cis-*inhibit Notch1 but *cis-*activate Notch2 signaling, while both Jag1 and Jag2 only *cis-*inhibit both Notch receptors(93). A systematic examination of *cis-* and *trans-* interactions between Jag2 and different Notch receptors could yield more profound insights into cell-cell communication-mediated MuSC functions.

### Notch activity modifiers

Activation of Notch signaling induces MuSC proliferation, self-renewal, and maintenance, and suppresses terminal myogenic differentiation via the *trans*-activation of Notch effector genes belonging to the negative bHLH family, including *Hes1*, *Hes5*, *Hey1* and *HeyL*, by antagonizing MyoD activity(94, 95). Our data indicated that *Jag2* mutant MuSCs isolated from homozygous *Jag2^sm^* and MuSC-specific *Jag2* KO mice display myogenic differentiation defects. The *Jag2* deficiency-mediated reduced myogenic differentiation is due to the failed *cis-*inhibition of Notch activity, since overexpression of reference *JAG2* but not pathogenic variants suppresses Notch activity and promotes myogenic differentiation.

Several Notch modifying factors have been identified. The Notch-controlled ankyrin repeat molecule (NRARP) functions as a counteracting regulator of Notch activity in many cell types in a negative feedback loop(96, 97). Notch3 KO mice display increased quiescent MuSCs and muscle hypertrophy due to hyperplasia of MuSC-derived myogenic precursor cells during muscle regeneration, potentially by upregulation of NRARP to suppress Notch1 activity(98). Numb regulates asymmetrical cell division, with one daughter cell inheriting Numb and the other inheriting Notch via antagonizing Notch activity(99). Reduced Notch signaling through elevated Numb expression in MuSCs resulted in myogenic differentiation by suppressing Notch activity(100, 101). Prox1-expressing MuSCs undergo myogenic differentiation through reciprocal suppression of Notch1(102). The Fringe homologs Lunatic fringe (Lfng), Manic fringe (Mfng), and Radical fringe (Rfng) are β3-N-acetylglucosaminyltransferases that influence Notch function by altering O-fucose modifications on EGF repeats in Notch receptors. Lfng enhances Notch2 activation via Dll1 and Dll4, whereas Mfng suppresses Notch2 activation through Jag1 and Jag2(103). Lfng enhances Dll1/4-driven *trans-* and *cis-*activation for Notch receptors(93). Thus, cellular environments with different expression levels of Notch receptors and Notch modifiers may influence Jag2-mediated Notch signaling.

### Notch for muscular dystrophies

Reduced MuSC self-renewal, maintenance, and differentiation contribute to the disease mechanism of muscular dystrophies. Therapies targeting Notch signaling could selectively enhance MuSC replication, potentially alleviating the symptoms of muscular dystrophy. Two other inherited muscle disease genes aside from *JAG2* have been linked to the Notch signaling pathway: *MEGF10* (104, 105) and *POGLUT1* (106). We determined that Megf10 and Notch1 interact at their intracellular domains and that this interaction is impaired by pathogenic variants(107). Loss of Notch signaling in *POGLUT1* deficiency has been demonstrated in skeletal muscle tissue from affected individuals and in *Drosophila*(106). *Jag1*-mediated augmentation of Notch signaling ameliorated DMD in canine and zebrafish models(77).

### Conclusions

We have demonstrated that Jag2-mediated *trans-*activation and *cis-*inhibition of Notch signaling regulate muscle stem cell function during muscle regeneration. JAG2 shows promise as a therapeutic target for muscular dystrophy, and our findings will help fine-tune interventions to focus on specific desirable downstream effects of JAG2-related interventions.

## METHODS

### Mice

C57BL/6N-*A^tm1Brd^ Jag2^tm1a(KOMP)Wtsi^*/HMmucd (*Jag2^LoxP/LoxP^*; MMRRC stock # 048257-UCD) were obtained from the Mutant Mouse Resource & Research Centers (MMRRC). *B6.Cg-Pax7^tm1(cre/ERT2)Gaka/J^* [*Pax7^+/CreERT2^*; JAX stock# 017763 (65)], *B6.Cg-Gt(ROSA)^26Sortm9(CAG-tdTomato)Hze/J^* [*Ai9*; JAX stock# 007909 (108)], and *STOCK Jag2^sm^/J* [*Jag2^sm^*; JAX stock# 000239;(80)], and B6.129S4-Gt(ROSA)26Sor^tm2(FLP*Sor^/J (*FLP*; JAX stock# 012930) were obtained from Jackson Laboratory. *Kdr^tm2.1Jrt/J^* (*Flk1^+/GFP^*) were obtained from Masatsugu Ema (109). *Cdh5^+/CreERT2^* mice were obtained from Dr. Yoshiaki Kubota (110). *B6.Cg-Pax7^tm1(cre/ERT2)Gaka/J^*(*Pax7^+/CreERT2^*) mice were crossed with *B6.Cg-Gt(ROSA)^26Sortm9(CAG-tdTomato)Hze/J^* (*Ai9*) to yield the *Pax7^+/CreERT2^:R26R^tdT^*(*Pax7^tdT^*) mice. *Pax7^tdT^* mice were bred with *Jag2^LoxP/LoxP^*and *Flk1^+/GFP^* to yield *Jag2^LoxP/LoxP^:Pax7^+/tdT^: Flk1^+/GFP^* mice. *Cdh5^+/CreERT2^* mice were crossed with *B6.Cg-Gt(ROSA)^26Sortm9(CAG-tdTomato)Hze/J^*(*Ai9*) to yield *Cdh5^+/CreERT2^:R26R^tdT^*(*Cdh5^tdT^*) mice. *Cdh5^tdT^* mice were bred with *Jag2^LoxP/LoxP^* and *Flk1^+/GFP^* to yield *Jag2^LoxP/LoxP^:Cdh5^+/tdT^: Flk1^+/GFP^* mice. *Jag2^sm^* mice(80) were crossed with *Pax7^tdT^*to yield the *Jag2^sm^:Pax7^tdT^* mice. All mouse colonies were established (Table 1) and genotyped (Table S6) in the laboratory. Cre recombination was induced using tamoxifen (TMX; T5648, MilliporeSigma) 75mg/kg body weight x 3 over 1 week at 3–6 weeks of age. *CreERT2* mice were used as controls. TA muscle regeneration was induced by intramuscular injection of 20µl of 10µM cardiotoxin (CTX) (V9125, MilliporeSigma). The animals were housed in an SPF environment and were monitored by Research Animal Resources (RAR) of the University of Minnesota. All protocols (2204–39969A) were approved by the Institutional Animal Care and Usage Committee (IACUC) of the University of Minnesota and complied with NIH guidelines for the use of animals in research.

### Cell culture and immunostaining

C2C12 myoblasts (CRL-1772) were obtained from American Type Culture Collection (ATCC) and cultured in DMEM medium with 10% FBS, 100units/ml of penicillin, and 100μg of streptomycin at 37°C in 5% O_2_ and 5% CO_2_. C2C12 cells were STR profiled to confirm their identity and tested negative for mycoplasma. MuSCs were isolated from adult mice(111). After collagenase type II (CLS-2, Worthington) treatment, dissociated cells from mouse hindlimb muscles were incubated with anti-CD31-PE (12-0311-82, eBiosciences), anti-CD45-PE (12-0451-81, eBiosciences), anti-Sca1-PE (A18486, eBiosciences), and anti-Integrin α7 (ABIN487462, MBL International), followed by anti-PE microbeads (130-048-801, Miltenyi Biotec), then underwent LD column (130-042-901, Miltenyi Biotec) separation. Negative cell populations were incubated with anti-Mouse IgG beads (130-048-402, Miltenyi Biotec), and then MS column (130-042-201, Miltenyi Biotec) separation was performed to isolate Integrin α7(+) MuSCs. MuSCs were maintained in culture on collagen-coated plates in myoblast growth medium containing 20% FBS, 20ng/ml bFGF (PHG0263, Invitrogen), 100units/ml of penicillin, and 100μg of streptomycin in HAM’s-F10 medium. MuECs were isolated from adult mice(14). Dissociated muscle cells were obtained as described above. Dissociated cells were incubated with CD45 MicroBeads (130-052-301, eBiosciences) and anti-CD45-PE (12-0451-81, eBiosciences), and then underwent LD column (130-042-901, Miltenyi Biotec) separation. Negative cell populations were incubated with CD31 MicroBeads (130-097-418, Miltenyi Biotec), and then MS column (130-042-201, Miltenyi Biotec) separation was performed to isolate CD45(-)CD31(+) MuECs. MuECs were maintained in culture on fibronectin-coated plates in EGM-2 Endothelial Cell Growth Medium-2 Bullet Kit (CC-3162, Lonza). All antibody-related materials are listed in Table S7. All cell cultures were maintained in a humidified incubator at 37°C with 5% CO_2_ and 5% O_2_.

4-Hydroxy tamoxifen (4-OHT, H6278, MilliporeSigma) treatment (1µM in EtOH) was used to induce *Jag2* deletion in MuSCs isolated from *Jag2^LoxP/LoxP^:Pax7^CreERT2^* mice. For the cell proliferation assay, cells were exposed to 1µM EdU for 3 hours before being fixed and stained by the Click-iT EdU Alexa Fluor 488 or 647 Imaging Kit (C10337 or C10340, Fisher Scientific). To induce differentiation of MuSCs, the myoblast growth medium was replaced with differentiation medium that contained DMEM supplemented with 5% horse serum for 1 to 5 days. Following cell cultures, anti-MyHC (MF20, DSHB) and anti-MyoD (C-20, Santa Cruz Biotechnology) antibodies followed by secondary anti-mouse-Alexa-488 (A11001, Thermofisher Scientific) and anti-rabbit-Alexa-647 (A32795, Thermofisher Scientific), anti-MyHC (MF20, DSHB) and anti-Hes1 (D6P2U, Cell Signaling Technology) antibodies followed by secondary anti-mouse-Alexa-568 (A11004, Thermofisher Scientific) and anti-rabbit-Alexa-488 (A11008, Thermofisher Scientific), or anti-Jag2 antibody (C23D2, Cell Signaling Technology) followed by secondary anti-rabbit-Alexa-488 (A11008, Thermofisher Scientific) were used (Table S7). LacZ expression in MuSCs obtained from *LacZ/Neo-Jag2^LoxP/LoxP^* mice was detected by X-gal staining overnight as described previously (15). Following X-gal staining, anti-Pax7 (DSHB) or anti-MyoD (C-20, Santa Cruz Biotechnology) antibodies followed by secondary anti-mouse-Alexa-488 (#A-11001, Thermofisher Scientific) or anti-rabbit-Alexa-488 (#A-11008, Thermofisher Scientific) were used. The fusion index (containing two or more nuclei in MyHC-positive myotubes) was measured. For induction of apoptotic cell death in MuSCs, Thapsigargin-mediated apoptosis was induced by 1µM of thapsigargin (T9033, MilliporeSigma) dissolved in EtOH for 24 hours. UV light-mediated apoptosis was induced by exposing the cells to UV light in a cell culture hood for 1 min without medium. After UV exposure, cell survival was assessed 24 hours following culture in 0.1% FBS in HAM’s F10 medium using the Annexin V Assay Kit (ab232855, Abcam). DAPI was used for nuclei staining.

### shRNA knockdown of *Jag2* in C2C12 mouse myoblasts

C2C12 cells were transfected with a cocktail of shRNA plasmids (Genecopoeia) against mouse *Jag2* using Lipofectamine 3000 (L3000001, ThermoFischer Scientific). The *Jag2* and scrambled shRNA sequences are shown in Table S6. shRNA transfection and positive clone selection were performed as described(30).

### Notch plate array and myotube-fusion index of differentiated C2C12 cells

Scrambled and *Jag2* shRNA cells were switched at 80% confluence from normal growth medium containing 10% FBS to differentiation medium containing 2% horse serum and differentiated for 4 days. Total RNA was isolated using an RNA isolation kit (Zymo Research). Reverse transcription of mRNA was performed using a high-capacity RNA to cDNA kit (Applied Biosystems). qPCR-based gene expression analysis was conducted using the TaqMan Fast Advanced Master Mix in the QuantStudio 3 Real-Time PCR System (ThermoFisher Scientific). The Taqman probes used were: mouse *Jag2* (Mm01325629_m1); mouse *MyoG* (Mm00446194_m1); human *JAG2* (Hs99999198_m1), and mouse *Gapdh* (Mm99999915_g1) from ThermoFisher Scientific. The cDNA samples were also examined via the TaqMan Array Mouse Notch Signaling Pathway, Fast 96-well plate (ThermoFisher Scientific) containing a set of Notch signaling pathway-associated genes and endogenous control genes as reported(30). For myotube fusion index analysis, the cells were fixed in 4% paraformaldehyde (PFA) for 15 mins and blocked in serum medium for 1 hour. The cells were stained with MHC [MF20, Developmental Study Hybridoma Bank (DSHB)] primary antibody for 1 hour. Cells were then stained with anti-rabbit or anti-mouse Alexa Fluor-568 secondary antibody for 1 hour. Nuclei were stained using DAPI (4’,6-diamidino-2-phenylindole dihydrochloride, D1306, Thermofisher Scientific), and the coverslips were mounted using Fluoromount Aqueous Mounting Medium (MilliporeSigma). The slides were imaged using a DM6000B or DM5500B epifluorescent microscope (Leica). The myotube-fusion index was determined by counting the number of nuclei within MHC-positive myotubes divided by the total number of nuclei in the field of view using ImageJ (National Institutes of Health).

### Site-directed Mutagenesis and overexpression of *JAG2* variants in C2C12 myoblasts

Site-directed mutagenesis was performed on the *JAG2* coding sequence in the *pCDNA3.1* backbone to generate the variants of interest using the Q5 Site-Directed Mutagenesis Kit (New England Biolabs). *pCDNA3.1JAG2* was a gift from Sandra Coppens. Mutagenic primers harboring the desired variants were used in a PCR reaction with wild-type *JAG2* sequence as the template. PCR was performed using the primers for the indicated *JAG2* variants (*p.Glu164Lys*, *p.Pro682Ser*, and *p.Phe977Ser*) (Table S6). Generation and the stable overexpression of the *JAG2* human variants in scrambled and *Jag2* shRNA cells was performed.

### RNA and genomic DNA isolation and qPCR

Cultured cells were washed with ice-cold PBS and lysed in place with Trizol. RNA was isolated using the DirectZol RNA Microprep Kit (R2062, Zymo Research) with on-column DNase digestion followed by cDNA synthesis using the Transcriptor First Strand cDNA synthesis kit (04379012001, Roche Molecular Diagnostics) with random primers. Genomic DNA for genotyping was isolated from mouse tail snips with lysis buffer containing Proteinase K (P2308, MilliporeSigma). qPCR was performed using GoTaq qPCR Master Mix (A6002, Promega). The input RNA amount was normalized across all samples, and *18S rRNA* was used for normalization of qPCR across samples. All primers were synthesized as custom DNA oligos from Integrated DNA Technologies (IDT) (Table S6).

### Single-muscle fiber isolation and staining

Extensor digitorum longus (EDL) muscle was dissected from uninjured or CTX-injected lower hindlimb muscle and digested with 0.2% collagenase type I (C0130, MilliporeSigma) for single muscle fiber isolation(15). Single muscle fibers were fixed with 2% PFA/PBS, permeabilized with 0.2% Triton-X100. Anti-Pax7 antibody (DSHB), followed by secondary anti-mouse-Alexa-488 (#A-11029, Thermofisher Scientific), was used for immunostaining (Table S7). DAPI was used for nuclear staining.

### Co-culture

Co-culture was performed by plating a monolayer of MuECs overlaid with MuSCs and culturing them in low-serum media to induce myogenic differentiation and reserve cell induction for 5 days (14). *Jag2* siRNAs (sc-39673, Santa Cruz Biotechnology) and control scramble siRNA-A (sc-37007, Santa Cruz Biotechnology) were transfected in ECs by Polyjet (11668019, ThermoFisher Scientific) before co-culture. A γ-secretase inhibitor, DAPT 10 μM, (D5942, Sigma-Aldrich), was used to block Notch signaling. Following co-cultures, anti-Pax7 (DSHB) and anti-MyoD (C-20, Santa Cruz Biotechnology) antibodies followed by secondary anti-mouse-Alexa-488 (#A-11029, Thermofisher Scientific) and anti-rabbit-Alexa-594 (#A-21207, Thermofisher Scientific) were used to detect Pax7(+)MyoD(-) self-renewing reserve cell population (Table S7). DAPI was used for nuclear staining.

### Luciferase reporter assays

The firefly luciferase reporter genes *Hes1-467-luciferase* [*pHes1*(*467*)*-Luc*] and *Hes1-467-Mut-luciferase* [*pHes1(467 RBPj(-)-Luc*] were obtained from Addgene [41723 and 43805, Addgene; (112)]. *4R-SV-luciferase* (*4R-SV-Lux*) was obtained from Andrew Lassar (67). *pRL-TK* (E1910, Promega) was used as an internal control. WT and homozygous MuSCs were transfected with expression vectors for human *JAG2* (*pcDNA3-JAG2*), mouse *MyoD* (*pcDNA3-MyoD*), mouse *Notch1* (*pCS2-Notch1*), mouse *Notch2* (*pPB[Exp]-Notch2*, VectorBuilder), human *NOTCH3* (*pPB[Exp]-NOTCH3*, VectorBuilder), mouse *Notch4* (*pHyTc-Notch4*, Addgene), or empty vectors, and the luciferase reporter genes using PolyJet™ In Vitro DNA Transfection Reagent (SL100688, SignaGen Laboratories). Cells were harvested 48 hours after transfection. Luciferase activity was measured with a plate reader (LD400; Beckman Coulter) using a dual luciferase reporter assay system (E1910, Promega).

### Ligand-coating Notch signaling assay

A total of 5×10^4^ WT MuSCs, *Jag2^sm^* homozygous MuSCs, WT MuSCs carrying *JAG2*, and *Jag2^sm^* homozygous MuSCs carrying *JAG2* were placed in a 48-well tissue culture plate pretreated with 5 μg/ml of DLL1-Fc, DLL4-Fc, JAG1-Fc, or JAG2-Fc (5026-DL, 10089-D4, 10969-JG, and 4748-JG, R&D) and allowed to settle at room temperature for 1 hour. After 16–18 hours, the cells were transfected with *pHes1*(*467*)*-Luc* using PolyJet™ In Vitro DNA Transfection Reagent (SL100688, SignaGen Laboratories). After 3 hours, the medium was changed to growth medium for an additional 48 hours. Cells were harvested for luciferase assays and RNA isolation.

### Histology and immunostaining for sections and cell cultures

The mouse tibialis anterior (TA) muscle was used for all histological analyses. Tissues were snap-frozen using LiN_2_ chilled isopentane and stored at –80°C. Eight-μm thick transverse cryosections were used for histological analysis. Hematoxylin & Eosin (H&E) staining was performed(15). Sirius red (Direct Red 80, 365548, MilliporeSigma) staining was performed on muscle sections to detect fibrosis(75). Muscle sections were stained in Oil Red O solution (O1391-250ML, MilliporeSigma)(76). LacZ expression in whole muscle and MuSCs obtained from *LacZ/Neo-Jag2^LoxP/LoxP^*mice was detected by X-gal staining overnight as described previously (15). Muscle sections obtained from X-gal-stained muscle were used for anti-CD31 antibody staining followed by anti-rat Alexa-488 (A11006, ThermoFisher Scientific). Anti-eMyHC (F1.652, DSHB) and anti-Laminin (L0663, MilliporeSigma) antibodies, followed by anti-mouse Alexa-488 (A11001, ThermoFisher Scientific) and anti-rat Alexa-568 antibodies (A11077, ThermoFisher Scientific) were used to detect regenerating muscle fibers. For capillary density measurement, anti-CD31 antibody (550274, BD Biosciences) and anti-Laminin antibody (L9393, MilliporeSigma) were used for TA sections, followed by anti-rat Alexa-488 (A11006, ThermoFisher Scientific) and anti-rabbit Alexa-568 (A11011, ThermoFisher Scientific). Immunostaining was performed on 35mm tissue culture plates or 8-well Permanox® Chamber slides (C7182-1PAK, MilliporeSigma). Cells were fixed with 2% PFA for 5 min and immunostained(15). Cells were permeabilized with 0.2% Triton-X in PBS, blocked with 1% BSA in PBS, and incubated with primary antibodies followed by secondary antibodies. PBS with 0.01% Triton-X was used for washing cells. Nuclei were counterstained with DAPI. The antibodies are listed in Table S7. Microscopic images were captured by a DP-1 digital camera attached to a BX51 fluorescence microscope with 10×, 20×, or 40×UPlanFLN objectives with cellSens Entry 1.11 (Olympus). Photoshop (Adobe) and Fiji (NIH) were used for image processing and enumerating Feret’s diameters(77).

### Over-expression experiments

The *pcDNA3* expression vectors for WT human *JAG2*, human *JAG2* pathogenic variants (*p.Glu164Lys*, *p.Pro682Ser*, and *p.Phe977Ser*), and mouse *MyoD* were transfected to low-passaged (2–3 passages) WT or homozygous *Jag2^sm^* MuSCs using PolyJet™ In Vitro DNA Transfection Reagent (SL100688, SignaGen Laboratories) for 3 hours. For the generation of the stable cell lines, the medium was replaced with myoblast growth medium, and G418 (Geneticin, 300 μg/ml; 10131035, Gibco) was added for the selection of transformant cells, which were used for cell growth and differentiation assays.

### Proximity ligation assay (PLA)

Cells were inoculated into 3-cm culture dishes. The following day, cells were fixed with 4% PFA, followed by permeabilization for 5 minutes using PBS + 0.2% Triton X-100. After rinsing, the PLA was carried out using a Duolink in situ PLA kit (DUO92101, MilliporeSigma). Following incubation with primary antibodies (anti-Jag2 and anti-Notch1 antibodies; Table S7), two distinct PLA probes (anti-rabbit minus and anti-mouse plus) were combined and incubated on the dishes for 1 hour. Microscopic images were captured as described above.

### Grip strength test

A forelimb grip strength test was performed(113). Mice were gently pulled by the tail after forelimb-grasping a metal bar attached to a force transducer (Grip Strength Meter, 1027CSM-D52, Columbus Instruments). Grip strength tests were performed by the same blinded examiner. Five consecutive grip strength tests were recorded, and then the mice were returned to the cage for a resting period of 20 min. Then, three series of pulls were performed, each followed by 20-min resting period. The average of the three highest values out of the 15 values collected was normalized to the body weight for comparison.

### Treadmill running

Exer-3/6 Treadmill (Columbus Instruments) was used for treadmill running tests(114). For acclimation, mice in each lane ran on a treadmill for 5 minutes at 10 m/min on a 0% uphill grade daily for 3 days. Then, the mice ran on a treadmill with a 10% uphill grade, starting at 10 m/min for 5 minutes. Then, every 2 minutes, the speed was increased by 2 m/min until exhaustion, defined as the mice’s inability to remain on the treadmill. The running time and distance were recorded.

### Rotarod test

Mice were trained on the rotarod (0890M-D54 Rotamex-5, Columbus Instruments) for 2 days before data collection(115). During each trial, mice were placed on the rod at 10 rpm for 60 seconds, and the rod accelerated from 10 to 30 rpm at 30-second intervals. The total maximum testing time was 240 seconds. Each trial was performed twice daily at 2-hour intervals for three consecutive days. The latency to fall was recorded, and the most prolonged latency was used for analysis.

### Transcriptome (RNA-seq) analysis

Three WT and 4 *Jag2^sm^* homozygous RNA samples were extracted from MuSCs isolated from hindlimb muscles. The RNA samples underwent Tapestation analysis (Agilent) to ensure RNA quality for long-read sequencing. 1μg of each RNA sample was used as input for cDNA synthesis library prep following the *Ligation sequencing V14 - Direct cDNA sequencing (SQK-LSK114)* protocol from Oxford Nanopore Technologies (ONT) until elution at the cDNA repair and end-prep step. The cDNA was eluted in 23.5μl water. Following cDNA end prep, barcodes were ligated so that the cDNA could be pooled. 22.5μl end-prepped cDNA, 2.5μl Native Barcode (NB05-11 from SQK-NBD114.24), and 25µl Blunt/TA Ligase Master Mix were combined, following the *Direct cDNA sequencing - native barcoding (SQK-DCS109 with EXP-NBD104 and EXP-NBD114)* protocol. 5μl EDTA was used to inactivate the ligation reaction after 20 min of incubation. Half of each barcoded cDNA sample was pooled into one library, and the remaining half was pooled into a second library. The rest of the library prep and loading followed the protocol for *Ligation sequencing amplicons - Native Barcoding Kit 24 V14 (SQK-NBD114.24)*. The barcoded pooled libraries were loaded on two different PromethION flow cells (FLO-PRO114M). Sequencing was performed on a P2-solo (ONT) for ∼23 hours, after data acquisition had plateaued. The raw data was base called using dorado/0.5.3 with the --min-qscore 7 --kit-name ’SQK-NBD114-24’ --no-trim flags, and the data were assembled to the GRCm39 mouse genome (GRCm39). The resulting .bam file was demultiplexed by barcode using dorado demux using the --emit-fastq flag. The resulting .fastq files from corresponding barcodes of the two different libraries were concatenated together, and then transcripts from each sample were counted using minimap2/2.17 htseq. The transcript counts were input into DESeq2 using R 4.3.0 using default analysis parameters, and significantly differentially expressed genes (padj < 0.05) were recorded. Genes with low expression (baseMean < 10) were filtered out of gene enrichment analysis, but the results from the whole transcriptome were saved. GO analysis was performed using Metascape v3.5.20250101(116). Gene information was obtained from NCBI database (https://www.ncbi.nlm.nih.gov/gene).=

### Transgenic Drosophila generation

Human reference *JAG2* and variants were cloned from the *pcDNA3* constructs into the *pUASTattB* vector using the EcoRI and XbaI sites for *Drosophila* expression via the in-fusion cloning method (Takara). The resulting plasmids were sequenced and verified (Eurofins Genomics). Transgenic animals were generated by Bestgene through φC31 integrase-mediated transgenesis on the second chromosome landing site, attP40.

### Drosophila husbandry

All *Drosophila* lines were maintained with standard Bloomington food at 25°C with 70% humidity and a 12-hour light:12-hour dark cycle. All experiments were conducted at 27°C to enhance transgene expression efficiency.

### Drosophila behavioral assays

For a flight assay, 25 flies from each group were collected and aged for four days. They were funneled into a 500-ml glass cylinder. The distribution of flies in the cylinder was recorded by a document camera (IPEVO), and the average scores from five independent experiments were calculated. For a negative geotaxis assay, 25 flies from each group were collected and aged for four days. Flies were tapped down by gently striking the vials on the surface of a table. Their climbing behaviors were recorded by a document camera (IPEVO) for 30 seconds. Movie clips were exported and analyzed by ImageJ. Statistical significance was calculated by a two-sided t-test.

### Statistical analysis

Statistical analysis was performed using Prism 10 (Graphpad). For comparison between two groups, an unpaired t-test was used. For comparison between multiple groups, a one-way ANOVA was used with multiple comparisons to the control. Distributions were compared using a chi-squared test. Graphing of the data was performed using Prism 10. All values are means ± SEM unless noted otherwise. *, p<0.05; **, p<0.01; ***, p<0.001.

## Acknowledgments

We thank the Minnesota Supercomputing Institute (MSI) and the University of Minnesota Imaging Center (UIC). We thank Dr. Yoshiaki Kubota for kindly providing *Cdh5^+/CreERT2^* mice. This work was supported by the Department of Defense (DoD) Award (HT9425-23-1-0461), Muscular Dystrophy Association (MDA) Research Grant (1297954) and Greg Marzolf Jr. Research Foundation to AA.

## Notes

### Competing Interest Statement

The authors have declared no competing interest.

## References

1. Chakkalakal JV, et al. The aged niche disrupts muscle stem cell quiescence. Nature. 2012;490(7420):355–360.

2. Chakkalakal JV, et al. Early forming label-retaining muscle stem cells require p27kip1 for maintenance of the primitive state. Development. 2014;141(8):1649–1659.

3. Schultz E. Satellite cell proliferative compartments in growing skeletal muscles. Dev Biol. 1996;175(1):84–94.

4. Shea KL, et al. Sprouty1 regulates reversible quiescence of a self-renewing adult muscle stem cell pool during regeneration. Cell Stem Cell. 2010;6(2):117–129.

5. Dumont NA, et al. Dystrophin expression in muscle stem cells regulates their polarity and asymmetric division. Nat Med. 2015;21(12):1455–1463.

6. Rhoads RP, et al. Satellite cell-mediated angiogenesis in vitro coincides with a functional hypoxia-inducible factor pathway. Am J Physiol Cell Physiol. 2009;296(6):C1321–1328.

7. Urciuolo A, et al. Collagen VI regulates satellite cell self-renewal and muscle regeneration. Nat Commun. 2013;4:1964.

8. Bentzinger CF, et al. Fibronectin regulates Wnt7a signaling and satellite cell expansion. Cell Stem Cell. 2013;12(1):75–87.

9. Bjornson CRR, et al. Notch signaling is necessary to maintain quiescence in adult muscle stem cells. Stem Cells. 2012;30(2):232–242.

10. Goel AJ, et al. Niche Cadherins Control the Quiescence-to-Activation Transition in Muscle Stem Cells. Cell Rep. 2017;21(8):2236–2250.

11. Abou-Khalil R, et al. Autocrine and paracrine angiopoietin 1/Tie-2 signaling promotes muscle satellite cell self-renewal. Cell Stem Cell. 2009;5(3):298–309.

12. Christov C, et al. Muscle satellite cells and endothelial cells: close neighbors and privileged partners. Mol Biol Cell. 2007;18(4):1397–1409.

13. Kostallari E, et al. Pericytes in the myovascular niche promote post-natal myofiber growth and satellite cell quiescence. Development. 2015;142(7):1242–1253.

14. Verma M, et al. Muscle Satellite Cell Cross-Talk with a Vascular Niche Maintains Quiescence via VEGF and Notch Signaling. Cell Stem Cell. 2018;23(4):530–543.e9.

15. Verma M, et al. Flt-1 haploinsufficiency ameliorates muscular dystrophy phenotype by developmentally increased vasculature in mdx mice. Hum Mol Genet. 2010;19(21):4145–4159.

16. Verma M, et al. Inhibition of FLT1 ameliorates muscular dystrophy phenotype by increased vasculature in a mouse model of Duchenne muscular dystrophy. PLoS Genet. 2019;15(12):e1008468.

17. Verma M, et al. Endothelial cell signature in muscle stem cells validated by VEGFA-FLT1-AKT1 axis promoting survival of muscle stem cell. Elife. 2024;13:e73592.

18. Chillakuri CR, et al. Notch receptor-ligand binding and activation: insights from molecular studies. Semin Cell Dev Biol. 2012;23(4):421–428.

19. Vargas-Franco D, et al. The Notch signaling pathway in skeletal muscle health and disease. Muscle Nerve. 2022;66(5):530–544.

20. Seandel M, et al. Generation of a functional and durable vascular niche by the adenoviral E4ORF1 gene. Proc Natl Acad Sci U S A. 2008;105(49):19288–19293.

21. Rios AC, et al. Neural crest regulates myogenesis through the transient activation of NOTCH. Nature. 2011;473(7348):532–535.

22. Mourikis P, et al. A critical requirement for notch signaling in maintenance of the quiescent skeletal muscle stem cell state. Stem Cells. 2012;30(2):243–252.

23. Bröhl D, et al. Colonization of the satellite cell niche by skeletal muscle progenitor cells depends on Notch signals. Dev Cell. 2012;23(3):469–481.

24. Morrison SJ, Spradling AC. Stem cells and niches: mechanisms that promote stem cell maintenance throughout life. Cell. 2008;132(4):598–611.

25. Butler JM, et al. Endothelial cells are essential for the self-renewal and repopulation of Notch-dependent hematopoietic stem cells. Cell Stem Cell. 2010;6(3):251–264.

26. Hadland BK, et al. Endothelium and NOTCH specify and amplify aorta-gonad-mesonephros-derived hematopoietic stem cells. J Clin Invest. 2015;125(5):2032–2045.

27. Kobayashi H, et al. Angiocrine factors from Akt-activated endothelial cells balance self-renewal and differentiation of haematopoietic stem cells. Nat Cell Biol. 2010;12(11):1046–1056.

28. Rafii S, et al. Human ESC-derived hemogenic endothelial cells undergo distinct waves of endothelial to hematopoietic transition. Blood. 2013;121(5):770–780.

29. Ottone C, et al. Direct cell-cell contact with the vascular niche maintains quiescent neural stem cells. Nat Cell Biol. 2014;16(11):1045–1056.

30. Coppens S, et al. A form of muscular dystrophy associated with pathogenic variants in JAG2. Am J Hum Genet. 2021;108(5):840–856.

31. Logan CV, et al. Mutations in MEGF10, a regulator of satellite cell myogenesis, cause early onset myopathy, areflexia, respiratory distress and dysphagia (EMARDD). Nat Genet. 2011;43(12):1189–1192.

32. Boyden SE, et al. Mutations in the satellite cell gene MEGF10 cause a recessive congenital myopathy with minicores. Neurogenetics. 2012;13(2):115–124.

33. Servián-Morilla E, et al. A POGLUT1 mutation causes a muscular dystrophy with reduced Notch signaling and satellite cell loss. EMBO Mol Med. 2016;8(11):1289–1309.

34. Ogasawara M, et al. CGG expansion in NOTCH2NLC is associated with oculopharyngodistal myopathy with neurological manifestations. Acta Neuropathol Commun. 2020;8(1):204.

35. Appella E, Weber IT, Blasi F. Structure and function of epidermal growth factor-like regions in proteins. FEBS Lett. 1988;231(1):1–4.

36. Vargas-Franco D, et al. The Notch signaling pathway in skeletal muscle health and disease. Muscle Nerve. [published online ahead of print: August 15, 2022]. 10.1002/mus.27684.

37. Shimizu K, et al. Binding of Delta1, Jagged1, and Jagged2 to Notch2 rapidly induces cleavage, nuclear translocation, and hyperphosphorylation of Notch2. Mol Cell Biol. 2000;20(18):6913–6922.

38. Shimizu K, et al. Physical interaction of Delta1, Jagged1, and Jagged2 with Notch1 and Notch3 receptors. Biochem Biophys Res Commun. 2000;276(1):385–389.

39. Fleming RJ, et al. The gene Serrate encodes a putative EGF-like transmembrane protein essential for proper ectodermal development in Drosophila melanogaster. Genes Dev. 1990;4(12A):2188–2201.

40. Shawber C, et al. Jagged2: a serrate-like gene expressed during rat embryogenesis. Dev Biol. 1996;180(1):370–376.

41. Valsecchi C, et al. JAGGED2: a putative Notch ligand expressed in the apical ectodermal ridge and in sites of epithelial-mesenchymal interactions. Mech Dev. 1997;69(1–2):203–207.

42. Luo B, et al. Isolation and functional analysis of a cDNA for human Jagged2, a gene encoding a ligand for the Notch1 receptor. Mol Cell Biol. 1997;17(10):6057–6067.

43. Deng Y, et al. Characterization, chromosomal localization, and the complete 30-kb DNA sequence of the human Jagged2 (JAG2) gene. Genomics. 2000;63(1):133–138.

44. Gray GE, et al. Human ligands of the Notch receptor. Am J Pathol. 1999;154(3):785–794.

45. Irvin DK, et al. Patterns of Jagged1, Jagged2, Delta-like 1 and Delta-like 3 expression during late embryonic and postnatal brain development suggest multiple functional roles in progenitors and differentiated cells. J Neurosci Res. 2004;75(3):330–343.

46. Stump G, et al. Notch1 and its ligands Delta-like and Jagged are expressed and active in distinct cell populations in the postnatal mouse brain. Mech Dev. 2002;114(1–2):153–159.

47. Sander GR, Powell BC. Expression of notch receptors and ligands in the adult gut. J Histochem Cytochem. 2004;52(4):509–516.

48. Sander GR, Brookes SJH, Powell BC. Expression of Notch1 and Jagged2 in the enteric nervous system. J Histochem Cytochem. 2003;51(7):969–972.

49. DeHart SL, Heikens MJ, Tsai S. Jagged2 promotes the development of natural killer cells and the establishment of functional natural killer cell lines. Blood. 2005;105(9):3521–3527.

50. Kared H, et al. Jagged2-expressing hematopoietic progenitors promote regulatory T cell expansion in the periphery through notch signaling. Immunity. 2006;25(5):823–834.

51. Johnson J, et al. Notch pathway genes are expressed in mammalian ovarian follicles. Mech Dev. 2001;109(2):355–361.

52. Vanorny DA, et al. Notch signaling regulates ovarian follicle formation and coordinates follicular growth. Mol Endocrinol. 2014;28(4):499–511.

53. Guo P, et al. Endothelial jagged-2 sustains hematopoietic stem and progenitor reconstitution after myelosuppression. J Clin Invest. 2017;127(12):4242–4256.

54. Palano MT, et al. Jagged Ligands Enhance the Pro-Angiogenic Activity of Multiple Myeloma Cells. Cancers (Basel). 2020;12(9):E2600.

55. Becam I, et al. A role of receptor Notch in ligand cis*-*inhibition in Drosophila. Curr Biol. 2010;20(6):554–560.

56. Fiuza U-M, et al. Mechanisms of ligand-mediated inhibition in Notch signaling activity in Drosophila. Dev Dyn. 2010;239(3):798–805.

57. Sprinzak D, et al. Cis*-*interactions between Notch and Delta generate mutually exclusive signalling states. Nature. 2010;465(7294):86–90.

58. Thambyrajah R, et al. Cis inhibition of NOTCH1 through JAGGED1 sustains embryonic hematopoietic stem cell fate. Nat Commun. 2024;15(1):1604.

59. Nandagopal N, Santat LA, Elowitz MB. Cis*-*activation in the Notch signaling pathway. Elife. 2019;8:e37880.

60. Gu JM, et al. An NF-κB - EphrinA5-Dependent Communication between NG2+ Interstitial Cells and Myoblasts Promotes Muscle Growth in Neonates. Developmental Cell. 2016;36(2):215– 224.

61. Vaz R, et al. Fibronectin promotes migration, alignment and fusion in an in vitro myoblast cell model. Cell and tissue research. 2012;348(3):569–78.

62. Bentzinger CF, et al. Article Fibronectin Regulates Wnt7a Signaling and Satellite Cell Expansion. Stem Cell. 2013;12(1):75–87.

63. Sidow A, et al. Serrate2 is disrupted in the mouse limb-development mutant syndactylism. Nature. 1997;389(6652):722–725.

64. Jiang R, et al. Defects in limb, craniofacial, and thymic development in Jagged2 mutant mice. Genes Dev. 1998;12(7):1046–1057.

65. Murphy MM, et al. Satellite cells, connective tissue fibroblasts and their interactions are crucial for muscle regeneration. Development. 2011;138(17):3625–3637.

66. Dong Z, et al. Gamma-Secretase Inhibitor (DAPT), a potential therapeutic target drug, caused neurotoxicity in planarian regeneration by inhibiting Notch signaling pathway. Sci Total Environ. 2021;781:146735.

67. Weintraub H, et al. MyoD binds cooperatively to two sites in a target enhancer sequence: occupancy of two sites is required for activation. Proc Natl Acad Sci U S A. 1990;87(15):5623– 5627.

68. Watanabe S, et al. MyoD gene suppression by Oct4 is required for reprogramming in myoblasts to produce induced pluripotent stem cells. Stem Cells. 2011;29(3):505–516.

69. Gunage RD, Reichert H, VijayRaghavan K. Identification of a new stem cell population that generates Drosophila flight muscles. eLife. 2014;3:e03126.

70. Deng M, et al. Single cell transcriptomic landscapes of pattern formation, proliferation and growth in Drosophila wing imaginal discs. Development. 2019;146(18):dev179754.

71. Zeng C, et al. Delta and Serrate are redundant Notch ligands required for asymmetric cell divisions within the Drosophila sensory organ lineage. Genes Dev. 1998;12(8):1086–1091.

72. de Celis JF, et al. Notch signalling mediates segmentation of the Drosophila leg. Development. 1998;125(23):4617–4626.

73. Dofash L, et al. Three novel missense variants in two families with JAG2-associated limb-girdle muscular dystrophy. Neuromuscul Disord. 2024;42:36–42.

74. Nikitin S, et al. Case Report: Exploring the clinical spectrum of LGMD R27: insights from a case study with homozygous pathogenic variant in the JAG2 gene. Front Pediatr. 2024;12:1414465.

75. Oda T, et al. Mutations in the human Jagged1 gene are responsible for Alagille syndrome. Nat Genet. 1997;16(3):235–242.

76. Li L, et al. Alagille syndrome is caused by mutations in human Jagged1, which encodes a ligand for Notch1. Nat Genet. 1997;16(3):243–251.

77. Vieira NM, et al. Jagged 1 Rescues the Duchenne Muscular Dystrophy Phenotype. Cell. 2015;163(5):1204–1213.

78. De Souza Leite F, et al. Muscle-specific increased expression of *JAG1* improves skeletal muscle phenotype in dystrophin-deficient mice [preprint]. 2025. 10.1101/2025.03.12.642857.

79. Lan Y, et al. The Jagged2 gene maps to chromosome 12 and is a candidate for the lgl and sm mutations. Mamm Genome. 1997;8(11):875–876.

80. Casey LM, et al. Jag2-Notch1 signaling regulates oral epithelial differentiation and palate development. Dev Dyn. 2006;235(7):1830–1844.

81. Matsakas A, et al. Muscle ERRγ mitigates Duchenne muscular dystrophy via metabolic and angiogenic reprogramming. FASEB J. 2013;27(10):4004–4016.

82. Fujimaki S, et al. The endothelial Dll4–muscular Notch2 axis regulates skeletal muscle mass. Nat Metab. 2022;4(2):180–189.

83. Fujimaki S, et al. Notch1 and Notch2 Coordinately Regulate Stem Cell Function in the Quiescent and Activated States of Muscle Satellite Cells. Stem Cells. 2018;36(2):278–285.

84. Low S, et al. Delta-Like 4 Activates Notch 3 to Regulate Self-Renewal in Skeletal Muscle Stem Cells. Stem Cells. 2018;36(3):458–466.

85. Eliazer S, et al. Heterogeneous levels of delta-like 4 within a multinucleated niche cell maintains muscle stem cell diversity. Elife. 2022;11:e68180.

86. Bi P, et al. Stage-specific effects of Notch activation during skeletal myogenesis. Elife. 2016;5:e17355.

87. Li H, et al. Fly Cell Atlas: A single-nucleus transcriptomic atlas of the adult fruit fly. Science. 2022;375(6584):eabk2432.

88. del Álamo D, Rouault H, Schweisguth F. Mechanism and significance of cis-inhibition in Notch signalling. Curr Biol. 2011;21(1):R40–47.

89. Yaron A, Sprinzak D. The cis side of juxtacrine signaling: a new role in the development of the nervous system. Trends Neurosci. 2012;35(4):230–239.

90. Chen D, et al. A new model of Notch signalling: Control of Notch receptor cis-inhibition via Notch ligand dimers. PLoS Comput Biol. 2023;19(1):e1010169.

91. Xu X, et al. Jag1-Notch cis*-*interaction determines cell fate segregation in pancreatic development. Nat Commun. 2023;14(1):348.

92. Formosa-Jordan P, Ibañes M. Competition in notch signaling with cis enriches cell fate decisions. PLoS One. 2014;9(4):e95744.

93. Kuintzle R, Santat LA, Elowitz MB. Diversity in Notch ligand-receptor signaling interactions. Elife. 2025;12:RP91422.

94. Fukada S, et al. Molecular signature of quiescent satellite cells in adult skeletal muscle. Stem Cells. 2007;25(10):2448–2459.

95. Den Hartog L, Asakura A. Implications of notch signaling in duchenne muscular dystrophy. Front Physiol. 2022;13:984373.

96. Lamar E, et al. Nrarp is a novel intracellular component of the Notch signaling pathway. Genes Dev. 2001;15(15):1885–1899.

97. Jarrett SM, et al. Extension of the Notch intracellular domain ankyrin repeat stack by NRARP promotes feedback inhibition of Notch signaling. Sci Signal. 2019;12(606):eaay2369.

98. Kitamoto T, Hanaoka K. *Notch3* Null Mutation in Mice Causes Muscle Hyperplasia by Repetitive Muscle Regeneration. Stem Cells. 2010;28(12):2205–2216.

99. Roegiers F, Jan YN. Asymmetric cell division. Curr Opin Cell Biol. 2004;16(2):195–205.

100. Conboy IM, Rando TA. The regulation of Notch signaling controls satellite cell activation and cell fate determination in postnatal myogenesis. Dev Cell. 2002;3(3):397–409.

101. Shinin V, et al. Asymmetric division and cosegregation of template DNA strands in adult muscle satellite cells. Nat Cell Biol. 2006;8(7):677–687.

102. Kivelä R, et al. The transcription factor Prox1 is essential for satellite cell differentiation and muscle fibre-type regulation. Nat Commun. 2016;7:13124.

103. Kakuda S, et al. Canonical Notch ligands and Fringes have distinct effects on NOTCH1 and NOTCH2. J Biol Chem. 2020;295(43):14710–14722.

104. Logan CV, et al. Mutations in MEGF10, a regulator of satellite cell myogenesis, cause early onset myopathy, areflexia, respiratory distress and dysphagia (EMARDD). Nat Genet. 2011;43(12):1189–1192.

105. Boyden SE, et al. Mutations in the satellite cell gene MEGF10 cause a recessive congenital myopathy with minicores. Neurogenetics. 2012;13(2):115–124.

106. Servián-Morilla E, et al. A POGLUT1 mutation causes a muscular dystrophy with reduced Notch signaling and satellite cell loss. EMBO Mol Med. 2016;8(11):1289–1309.

107. Saha M, et al. Consequences of MEGF10 deficiency on myoblast function and Notch1 interactions. Hum Mol Genet. 2017;26(15):2984–3000.

108. Madisen L, et al. A robust and high-throughput Cre reporting and characterization system for the whole mouse brain. Nat Neurosci. 2010;13(1):133–140.

109. Ema M, Takahashi S, Rossant J. Deletion of the selection cassette, but not cis*-*acting elements, in targeted Flk1-lacZ allele reveals Flk1 expression in multipotent mesodermal progenitors. Blood. 2006;107(1):111–117.

110. Okabe K, et al. Neurons limit angiogenesis by titrating VEGF in retina. Cell. 2014;159(3):584–596.

111. Motohashi N, Asakura Y, Asakura A. Isolation, culture, and transplantation of muscle satellite cells. J Vis Exp. [published online ahead of print: April 8, 2014];(86). 10.3791/50846.

112. Nishimura M, et al. Structure, chromosomal locus, and promoter of mouse Hes2 gene, a homologue of Drosophila hairy and Enhancer of split. Genomics. 1998;49(1):69–75.

113. Aartsma-Rus A, van Putten M. Assessing functional performance in the mdx mouse model. J Vis Exp. 2014;(85):51303.

114. Fukada S, et al. Genetic Background Affects Properties of Satellite Cells and mdx Phenotypes. The American Journal of Pathology. 2010;176(5):2414–2424.

115. Jin Q, et al. A GDF11/myostatin inhibitor, GDF11 propeptide-Fc, increases skeletal muscle mass and improves muscle strength in dystrophic mdx mice. Skeletal Muscle. 2019;9(1):16.

116. Zhou Y, et al. Metascape provides a biologist-oriented resource for the analysis of systems-level datasets. Nat Commun. 2019;10(1):1523.

