## Supplemental Figures for "*JAG2*-related muscular dystrophy and Notch signaling dysfunction in muscle stem cells"

### Figure S1

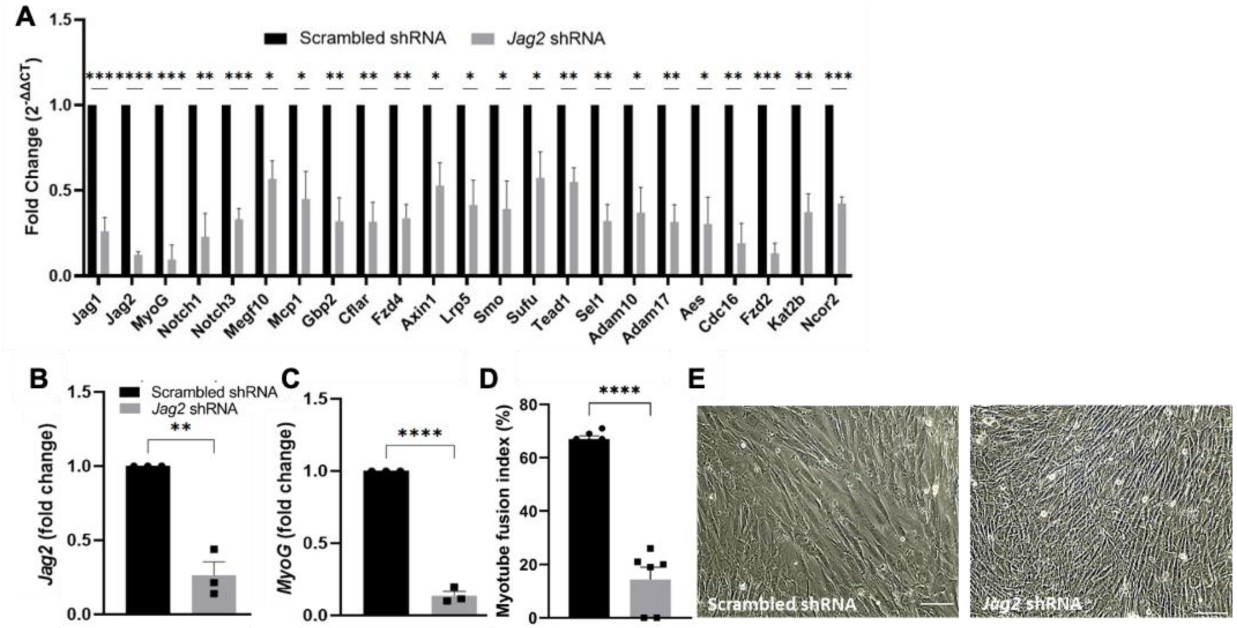


**Figure S1.**

**Impaired myogenic differentiation and Notch signaling in *JAG2* deficient-C2C12 myoblasts.** (A) *Jag2* shRNA and scrambled shRNA C2C12 myoblasts were differentiated for 6 days. qPCR was performed on isolated RNA with an array plate containing probes for 93 Notch signaling and myogenesis genes, along with 3 control genes. The 23 significantly downregulated genes are shown in the graph, including *MyoG*, *Notch1*, and *Notch3*. No genes were significantly upregulated. Transcript levels were normalized to *Gapdh*. *Jag2* shRNA and scrambled shRNA C2C12 myoblasts were differentiated for 4 days. RNA was isolated, followed by qPCR. (B) *Jag2* knockdown was confirmed. (C) *Jag2* shRNA knockdown cells showed lower levels of the late differentiation marker *MyoG* compared to controls. (D) *Jag2* knockdown cells show impaired myotube formation compared to scrambled shRNA controls. (E) Representative phase contrast images of differentiated *Jag2* knockdown cells show a lack of multinucleated myotube formation compared to scrambled shRNA cells. Scale bar, 100*μm*. n=3 independent experiments. Unpaired t-tests showed *, $p<0.05$; **, $p < 0.01$; ***, $p < 0.001$; ****, $p <0.0001$ for n=5 replicates. Error bars show the standard error of the mean (SEM).

### Figure S2

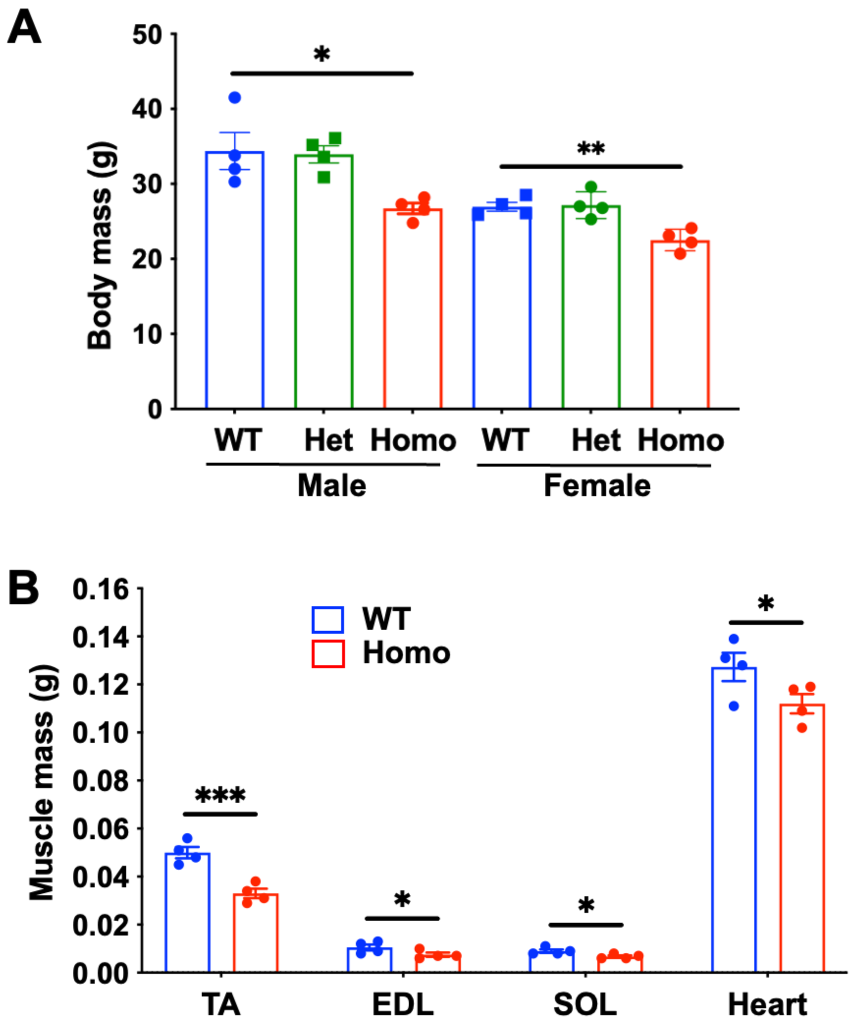


**Figure S2.**

**Reduced body and muscle mass in homozygous *Jag2*$^{sm}$ mice.** Three-month-old *Jag2*$^{sm}$ homozygous male and female mice showed reduced body mass, tibialis anterior (TA), extensor digitorum longus (EDL), soleus (SOL) muscle, and heart vs. *Jag2*$^{sm}$ WT and/or heterozygous mice. An unpaired t-test showed *, $p<0.05$; **, $p <0.01$. ***, $p <0.001$. Error bars show SEM.

### Figure S3

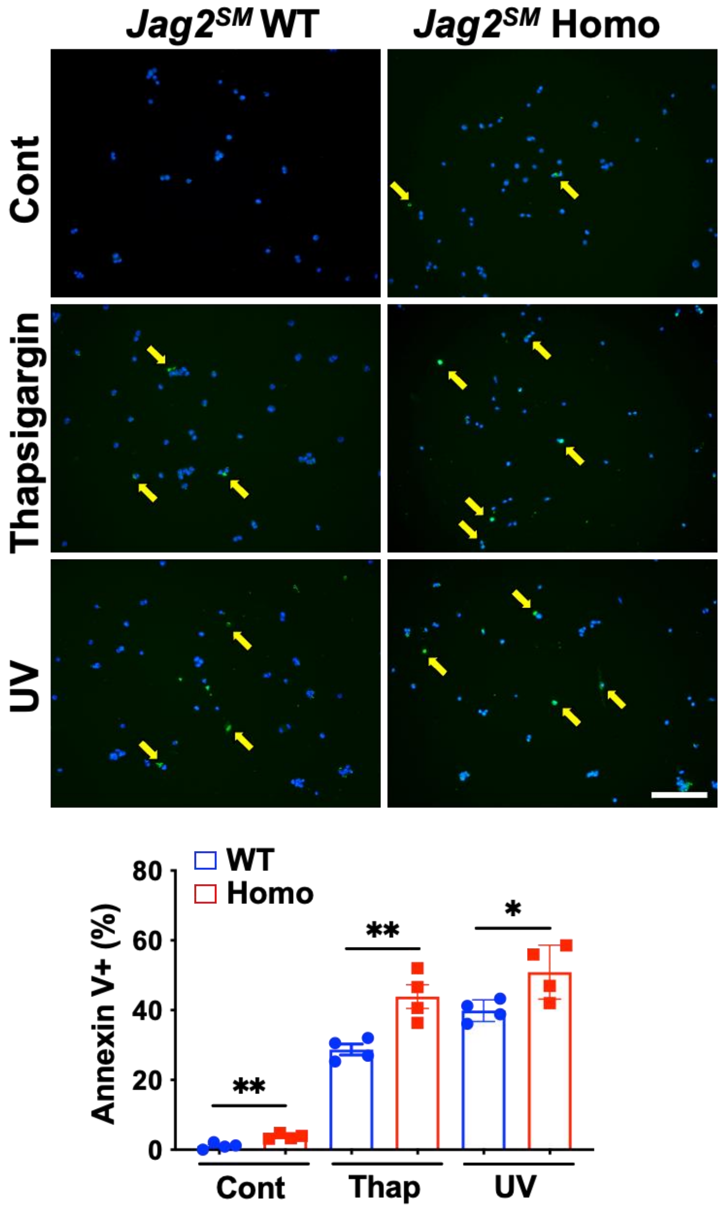


**Figure S3.**

***Jag2**$^{sm}$* **MuSCs show increased apoptotic cell death.** MuSCs isolated from WT and homozygous *Jag2*$^{sm}$ mice were used for apoptosis assays. Twenty-four hours following treatment with thapsigargin or UV exposure. Scale bars, 100 μm.

### Figure S4

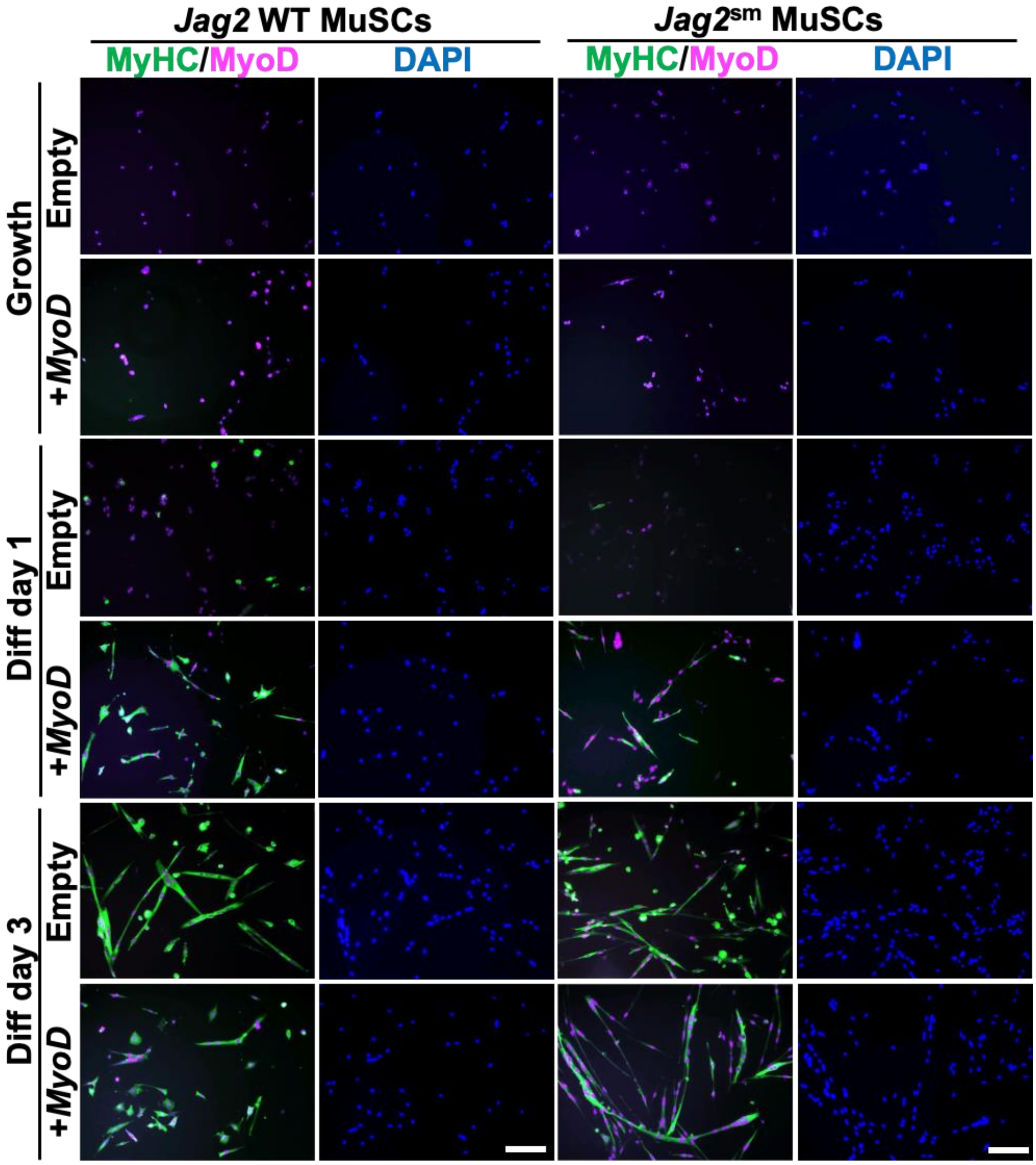


**Figure S4.**

**Overexpression of *MyoD* rescues differentiation defects in *Jag2$^{sm}$* MuSCs.** MuSCs isolated from WT and homozygous *Jag2$^{sm}$* mice were used for overexpression of *MyoD*. Overexpression of *MyoD* increased MyoD (purple) expression and MyHC(+) myogenic differentiation (green) and rescued myogenic differentiation defects in *Jag2$^{sm}$* MuSCs. Scale bars, 100 μm.

### Figure S5

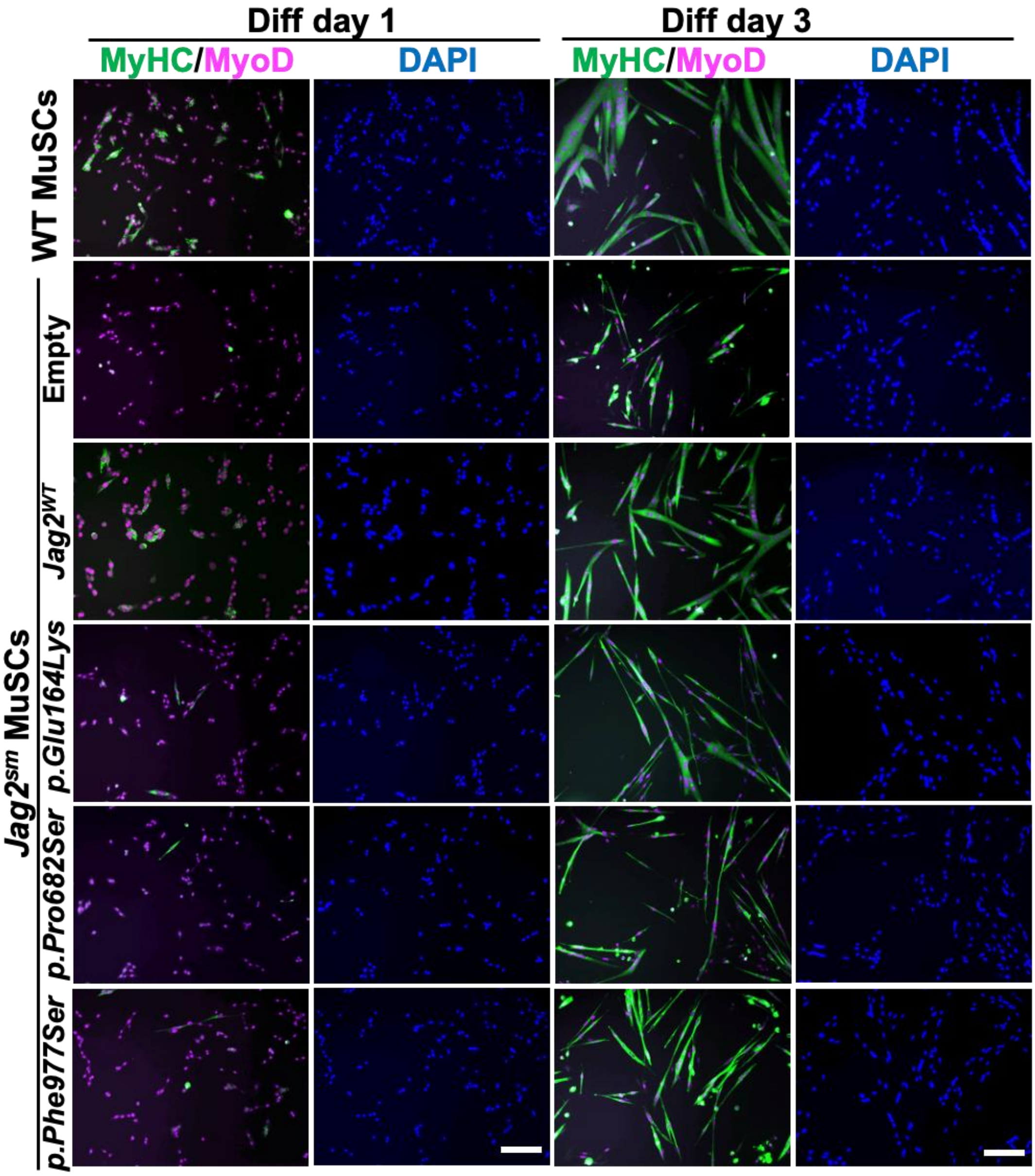


**Figure S5.**

**Overexpression of *JAG2* rescues differentiation defects in *Jag2*$^{sm}$ MuSCs.** MuSCs isolated from WT and homozygous *Jag2*$^{sm}$ mice were used for overexpression of human reference *JAG2* (*Jag2*$^{WT}$) and human *JAG2* harboring three patient variants (*p.Glu164Lys*, *p.Pro682Ser*, and *p.Phe977Ser*). Overexpression of *Jag2*$^{WT}$ *but not JAG2* variants increased expression and MyHC(+) myogenic differentiation (green) and rescued myogenic differentiation defects in *Jag2*$^{sm}$ MuSCs.. MyoD expression was shown in nuclei (purple). Scale bars, 100 µm.

### Figure S6

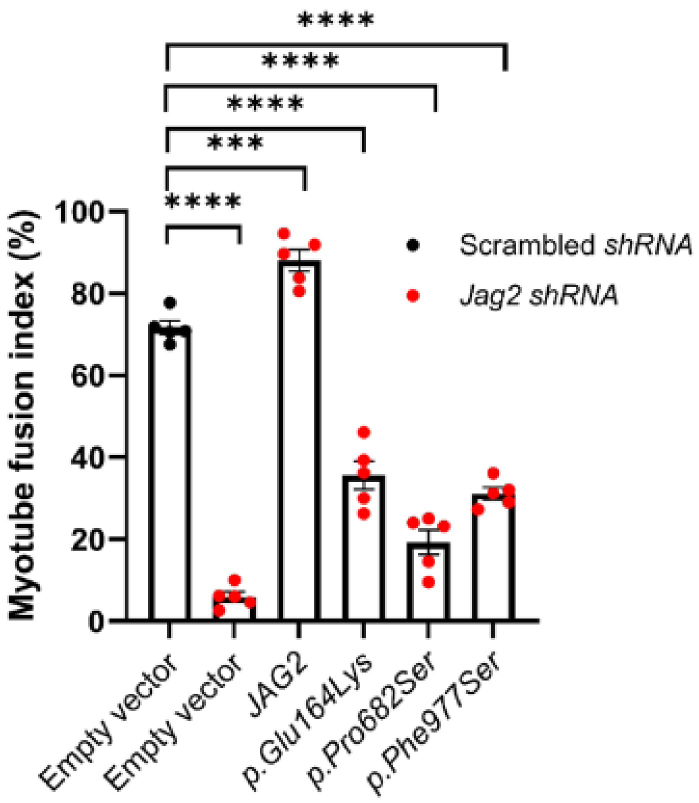


**Figure S6.**

**Rescue of *Jag2* shRNA cells**. *Jag2* shRNA cells with stable overexpression of empty vector, human reference *JAG2*, and human *JAG2* harboring three patient variants were differentiated for 6 days in 2% horse serum medium. Scrambled shRNA cells with stable overexpression of the empty vector were controls. The cells were fixed and stained with *MyHC* for quantification of the myotube-fusion index. Human reference *JAG2* overexpression rescued the effect of *Jag2* knockdown, whereas the variant forms of *JAG2* did not. An unpaired t-test showed ****, $p < 0.0001$; ***, $p < 0.001$. Error bars show the standard error of the mean (SEM).

### Figure S7

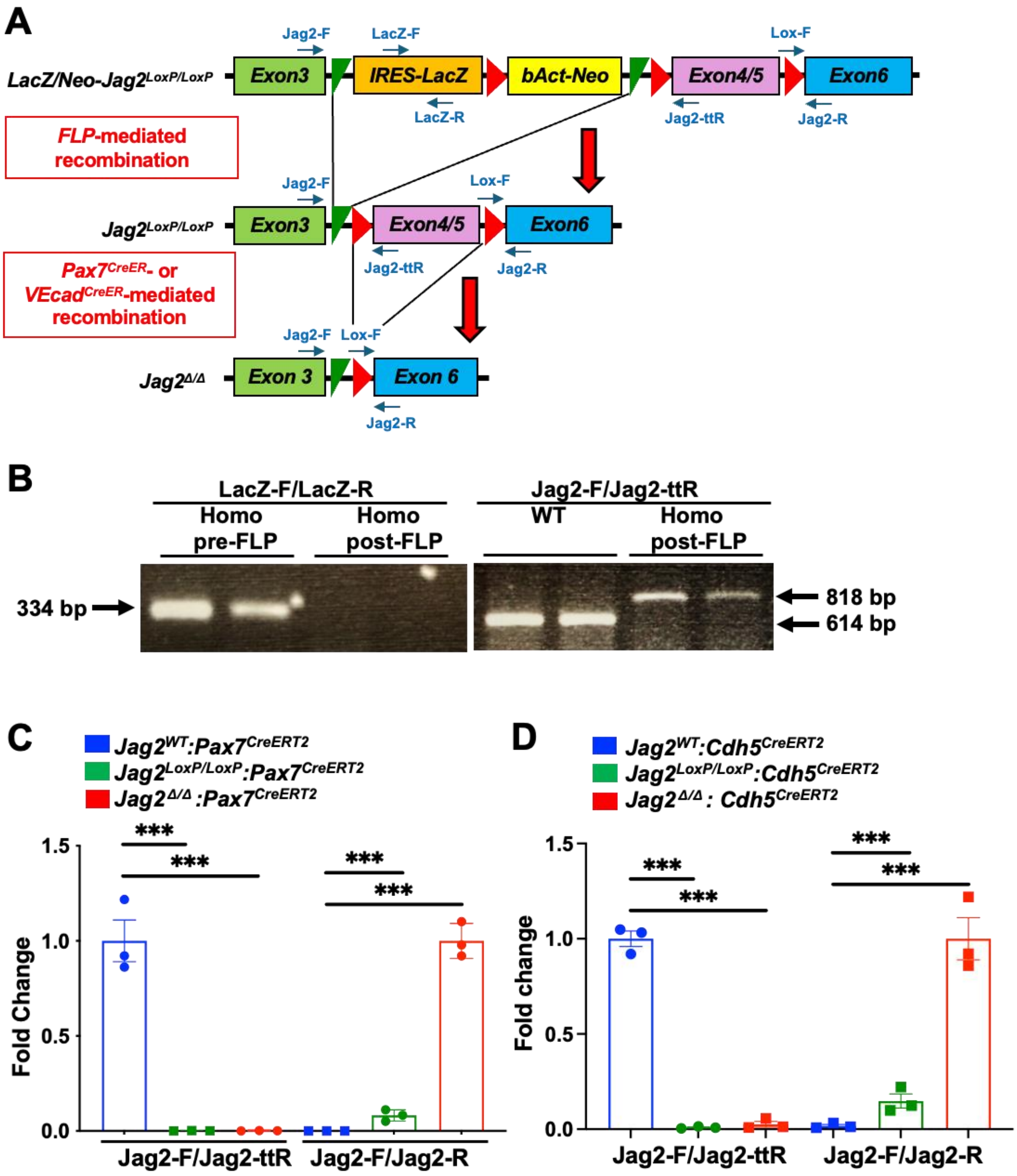
A
Jag2-F
LacZ-F
Lox-F
LacZ/Neo-Jag2LoxP/LoxP
Exon3
IRES-LacZ
bAct-Neo
Exon4/5
Exon6
LacZ-R
Jag2-ttR
Jag2-R
FLP-mediated recombination
Jag2LoxP/LoxP
Pax7CreER- or VEcadCreER-mediated recombination
Jag2Δ/Δ
Exon 3
Exon 6
B
LacZ-F/LacZ-R
Homo pre-FLP
Homo post-FLP
Jag2-F/Jag2-ttR
WT
Homo post-FLP
334 bp
818 bp
614 bp
C
Jag2WT:Pax7CreERT2
Jag2LoxP/LoxP:Pax7CreERT2
Jag2Δ/Δ :Pax7CreERT2
Fold Change
1.5
1.0
0.5
0.0
***
Jag2-F/Jag2-ttR
Jag2-F/Jag2-R
D
Jag2WT:Cdh5CreERT2
Jag2LoxP/LoxP:Cdh5CreERT2
Jag2Δ/Δ : Cdh5CreERT2
Fold change

**Figure S7.**

**Confirmation of conditional deletion of *Jag2*.** (A) Schematic genomic structure of *Jag2$^{floxed}$* locus for conditional *Jag2* mutant (*LacZ/Neo-Jag2$^{LoxP/LoxP}$*) mice. Following breeding with *FLP* mice, Flippase-Frt sites (green triangles)-mediated *IRES-LacZ/bAct-Neo* cassette deletion occurred. Following breeding with *Pax7$^{CreER}$* or *VEcad$^{CreER}$* mice and TMX-treatment, *CreER-LoxP* sites (red triangles)-mediated *Exons 4/5* deletion occurred. (B) Genomic PCR was performed to detect the LacZ gene in pre-FLP but not in post-FLP mice in homozygous *LacZ/Neo-Jag2$^{LoxP/LoxP}$* mice using *Lac-F/LacZ-R* primers. Following breeding with *FLP* mice, an 818 bp band appeared in homozygous *LacZ/Neo-Jag2$^{LoxP/LoxP}$* mice while WT mice showed an 614 bp band using *Jag2-F/Jag2-ttR* primers. (C) MuSCs isolated from *Jag2$^{LoxP/LoxP}$:Pax7$^{CreER}$* (without TMX) or *Jag2$^{\Delta/\Delta}$:Pax7$^{CreER}$* (with TMX) mice showed no DNA amplification by qPCR using *Jag2-F/Jag2-ttR* primers, while MuSCs from *Jag2$^{WT}$:Pax7$^{CreER}$* mice showed the DNA amplification. Following treatment with TMX, qPCR using *Jag2-F/Jag2-R* primers amplified the DNA in MuSCs from *Jag2$^{\Delta/\Delta}$:Pax7$^{CreER}$* mice, while *Jag2$^{WT}$:Pax7$^{CreER}$* or *Jag2$^{LoxP/LoxP}$:Pax7$^{CreER}$* mice showed no DNA amplification. (D) MuECs from *Jag2$^{LoxP/LoxP}$:VEcad$^{CreER}$* (without TMX) or *Jag2 $^{\Delta/\Delta}$:VEcad$^{CreER}$* (with TMX) mice showed no DNA amplification by qPCR using *Jag2-F/Jag2-ttR* primers, while *Jag2$^{WT}$:VEcad$^{CreER}$* showed DNA amplification. Following treatment with TMX, qPCR using *Jag2-F/Jag2-R* primers amplified DNA in MuECs from *Jag2$^{\Delta/\Delta}$:VEcad$^{CreER}$* mice but not in *Jag2$^{WT}$:VEcad$^{CreER}$* or *Jag2$^{LoxP/LoxP}$:VEcad$^{CreER}$* mice. An unpaired t-test showed ***, $p<0.001$. Error bars show the standard error of the mean (SEM).
